# The usefulness of multi-parent multi-environment QTL analysis: an illustration in different NAM populations

**DOI:** 10.1101/2020.02.03.931626

**Authors:** Vincent Garin, Marcos Malosetti, Fred van Eeuwijk

## Abstract

Commonly QTL detection in multi-parent population (MPPs) data measured in multiple environments (ME) is done by a single environment analysis on phenotypic values ‘averaged’ across environments. This method can be useful to detect QTLs with a consistent effect across environments but it does not allow to estimate environment-specific QTL (QTLxE) effects. Running separate single environment analyses is a possibility to measure QTLxE effects but those analyses do not model the genetic covariance due to the use of the same genotype in different environments. In this paper, we propose methods to analyze MPP-ME QTL experiments using simultaneously the data from several environments and modelling the genotypic covariances. Using data from the EU-NAM and the US-NAM populations, we show that these methods allow to estimate the QTLxE effects and that they give a more precise description of the trait genetic architecture than separate within environment analyses. The MPP-ME models we propose can also be extended to integrate environmental indices (e.g. temperature, precipitation, etc.) to understand better the mechanisms behind the QTLxE effects. Therefore, our methodology allows to exploit the full potential of MPP-ME data: to estimate QTL effect variations a) within the MPP between sub-populations due to different genetic backgrounds; and b) between environments.

## I. INTRODUCTION

The use of multi-parent populations (MPPs) to investigate biological questions becomes progressively a regular practice in plant genetics and plant breeding. Different MPPs have been developed like the nested association mapping (NAM) populations (McMullen et al., 2009) or the multi-parent advanced generation inter-cross (MAGIC) populations (Cavanagh et al., 2008). The collections of crosses between a set of parents used in breeding programs can also be analyzed as MPPs (Würschum, 2012; Parisseaux and Bernardo, 2004). Different statistical approaches have been proposed to detect QTLs in NAM populations (Xavier et al., 2015), in MAGIC designs (Verbyla et al., 2014) or in MPPs composed of crosses (Jourjon et al., 2005).

The plant phenotype is the result of cumulative interactions between the genotype and the environment (Malosetti et al., 2013). Therefore, researchers have developed statistical procedures to detect QTLs taking the genotype by environment (GxE) interactions into consideration (Boer et al., 2007; Korte et al., 2012). Several MPPs have been tested in multiple environments (MPP-ME) (Buckler et al., 2009; Giraud et al., 2014; Saade et al., 2016) but only few studies have proposed a proper MPP GxE QTL detection methodology (Piepho and Pillen, 2004; Verbyla et al., 2014). Most of the researches average the phenotypic values by calculating adjusted means or predictions across the environments that represent an average phenotypic value (Giraud et al., 2014; Poland et al., 2011; Buckler et al., 2009). In other articles, people performed separate analyses in each environment (e.g. Saade et al. (2016)).

We consider that the main interest of an MPP-ME QTL experiment is to estimate genetic (QTL) variations at two levels: a) within the MPP between sub-populations due to different genetic background; and b) between environments. In previous researches, we developed a framework for QTL detection in MPPs composed of crosses between a set of parents with different assumptions about the QTL effects (Garin et al., 2017, 2018). The QTL effects were assumed to be more or less diverse/consistent in the different genetic backgrounds, which allowed to estimate QTL allelic variations within the MPP. In the present paper, we extended our methodology to the multi-environment context. We allowed the QTL effects to also vary between the environments to estimate QTL by environment (QTLxE) interactions. We further extended our models to integrate environmental information like temperature or precipitation to get a deeper understanding of the mechanisms behind the QTLxE effects. In the following sections, we present different methods for analyzing MPP-ME QTL experiments. We illustrate the usefulness of our methodology with examples coming from the EU-NAM and the US-NAM populations.

## II. MATERIAL AND METHODS

### Statistical methodology

In Table 1, we present a list of models for analyzing MPP-ME data. The first set of models allows to model phenotypic variation. The second category represents genotypic models used to detect QTLs. The last section presents four methods for QTL detection in MPP-ME experiments. The models and their components are described in the following sections assuming that the data come from two environments.

**Table 1:** Models table

| Code | Model |
| --- | --- |
| **Phenotypic models** |  |
| 1 | *Genotype BLUEs within environment*<br>$y_{icp} = \mu + des + G_{ic} + \epsilon_{icp}$<br>BLUEs vectors: $y_{E1}, y_{E2}$ |
| 2 | *Genotype BLUEs across environments*<br>$y_{icep} = \mu_e + des_{(e)} + G_{ic} + GE_{ice} + \epsilon_{icep}$<br>BLUEs vector: $y_{E(1,2)}$ |
| **Genotypic models** |  |
| *Single environment analysis* |  |
| 3 | $y_{ic} = \mu_c + G_{ic} + \epsilon_{ic}$ |
| 3.1 | $G_{ic} = \sum_q x_{iq}^T * \beta_q + g_{ic}$ |
| 3.2 | $\epsilon_{ic} \sim N(0, R)$ with $R = \bigoplus_{c=1}^{n_c} \sigma_{\epsilon_c}^2$ |
| 3.3 | $y = X_c \beta_c + X_Q \beta_Q + \epsilon$ |
| *Multi-environment analysis* |  |
| 4 | $y_{ice} = \mu_{ce} + G_{ice} + \epsilon_{ice}$ |
| 4.1 | $G_{ice} = \sum_q x_{iq}^T * \beta_{qe} + g_{ice}$ |
| 4.2 | $g_{ice} \sim N(0, \sigma_g^2)$ |
| 4.3 | $\epsilon_{ice} \sim N(0, R)$ with $R = \bigoplus_{c=1}^{n_c} \sigma_{\epsilon_{c(e)}}^2$ |
| 4.4 | $\begin{bmatrix} y_1 \\ y_2 \end{bmatrix} = \begin{bmatrix} X_c & 0 \\ 0 & X_c \end{bmatrix} \begin{bmatrix} \beta_{c1} \\ \beta_{c2} \end{bmatrix} + \begin{bmatrix} X_Q & 0 \\ 0 & X_Q \end{bmatrix} \begin{bmatrix} \beta_{Q1} \\ \beta_{Q2} \end{bmatrix} + g + \epsilon$ |
| **MPP-ME QTL detection methods** |  |
| *M1: MPP QTL analysis on BLUEs across environments* |  |
| M1 | $y_{E(1,2)} = X_c \beta_c + X_Q \beta_Q + \epsilon$ |
| *M2: MPP QTL analysis on BLUEs within environment* |  |
| M2 | $y_{E1} = X_c \beta_c + X_Q \beta_Q + \epsilon$<br>$y_{E2} = X_c \beta_c + X_Q \beta_Q + \epsilon$ |
| *M3: Two-stage MPP GxE QTL analysis* |  |
| M3 | $\begin{bmatrix} y_{E1} \\ y_{E2} \end{bmatrix} = \begin{bmatrix} X_c & 0 \\ 0 & X_c \end{bmatrix} \begin{bmatrix} \beta_{cE1} \\ \beta_{cE2} \end{bmatrix} + \begin{bmatrix} X_Q & 0 \\ 0 & X_Q \end{bmatrix} \begin{bmatrix} \beta_{QE1} \\ \beta_{QE2} \end{bmatrix} + g + \epsilon$ |
| *M4: One-stage MPP GxE QTL analysis* |  |
| M4 | $y_{icep} = \mu_{ce} + des_{(e)} + G_{i'(e)} + \sum_q x_{iq}^T * \beta_{qe} + g_{ice} + \epsilon_{icep}$ |

### Phenotypic models

Models 1 and 2 model phenotypic variations accounting for experimental design factors (*des*) such as replicates, incomplete blocks, row-columns, etc. In model 1, *y*_*icp*_ is the plot data phenotypic value of genotype *i* from cross *c. G*_*ic*_ is the genetic effect of genotype that can be separated into the tested genotypes and the check entries like in Boer et al. (2007). Finally, *ϵ*_*icp*_ is the plot error. Model 1 allows to calculate the genotype best linear unbiased estimates (BLUEs) treating *G*_*ic*_ as fixed. In model 1, we considered the data from each environment separately and calculated within environment genotype BLUEs (***y***_***E*1**_, ***y***_***E*2**_).

In model 2, we model the phenotypic variation of the two environments jointly. *y*_*icep*_ is the phenotypic plot measurement of genotype *i* from cross *c* in environment *e*. The intercept *µ*_*e*_ and the design term *des*_(*e*)_ become environment specific. The term *GE*_*ice*_ represents the genotype by environment interaction. In model 1 and 2, all terms were fitted as random except the genotype term *G*_*ic*_. The genotype BLUEs (***y***_***E*(1**,**2)**_) obtained with model 2 represent phenotypic values across environments.

### Genotypic models

Models 3 and 4 are MPP QTL detection models for single and multi-environment data, respectively. In model 3, *y*_*ic*_ is the phenotypic value of the *i*^*th*^ individual in cross *c. µ*_*c*_ is a cross intercept. The term *G*_*ic*_ (3.1) describes the genotypic effect. *G*_*ic*_ can be partitioned into a fixed QTL part 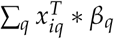 and a residual part 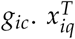 represents the expected number of QTL alleles received by individual *i* at position *q* and *β*_*q*_ are the QTL allelic substitution effects.

The vector 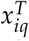 is of dimensions [1 × *n*_*al*_] and varies according to the number of alleles (*n*_*al*_) assumed at the QTL position. A first model called parental assumes that each parent contributes a unique allele to the MPP (*n*_*al*_ = *n*_*p*_). A second option called ancestral model assumes that genetically similar parents inherit their allele from a common ancestor. Parents are grouped in ancestral classes based on genetic similarity. We assume that each group represents one ancestral allele (*n*_*al*_ = *n*_*a*_). The final possibility is a bi-allelic model assuming that genotypes with the same SNP score transmit the same allele.

The elements of 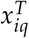 take values between 0 and 2. For the parental model, they represent the expected number of parental allele copies estimated using IBD probabilities computed by the package R/qtl (Broman et al., 2003). For the ancestral model the vector 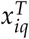 specifying the parental allele distribution is modified taking identical by state (IBS) parental genetic relatedness into consideration. We used the R package clusthaplo (Leroux et al., 2014) to evaluate the parent genetic similarities along the genome and infer common ancestral classes. For the bi-allelic model, 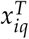 is a scalar taking values 0, 1 or 2 corresponding to the number of IBS copies of the minor SNP allele. The model is estimated fixing one QTL allele as reference. In NAM populations, it is convenient to set the central parent allele as reference and to interpret the additive allelic substitution effect *β*_*q*_ as the deviations with respect to the central parent.

*G*_*ic*_ (3.1) is also composed of a residual genetic effect *g*_*ic*_ that was not accounted by the QTLs. In the single environment analysis, *g*_*ic*_ is not directly modelled. The residual genetic variation is modelled by the error term *ϵ*_*ic*_ (3.2), which also contains the residual plot error. *ϵ*_*ij*_ follows a normal distribution *N*(0, ***R***). We assume that 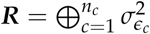 which means that the variance of the error term is different in each cross to take into consideration the heterogeneity that could exist between crosses.

Model 3 can be expressed in matrix notation (3.3) with ***y*** being the phenotypic values vector of dimension [*N* × 1]. ***X***_***c***_ is a [*N* × *n*_*c*_] cross-specific intercept matrix with ***β***_***c***_ representing the vectors of cross intercepts. ***X***_***Q***_ is a [*N* × *n*_*al*_] QTL incidence matrix and ***β***_***Q***_ the corresponding vector of QTL additive allelic substitution effects.

Model 4 is a modification of model 3 to analyze jointly the data from several environments. Compared to model 3, the cross effects *µ*_*ce*_, the QTL effects *β*_*qe*_ (4.1), and the residual genetic effect *g*_*ice*_ (4.2) are now indexed per environment. In expression 4.1, we modelled the environmental specific QTL allelic effects. These QTL allelic effects could also be partitioned into a main effect across environments and environment-specific components. This is a difference with respect to model 3. Another important difference between model 3 and 4, is the explicit modelling of the genotypic covariance between phenotypic observations measured on the same genotype in different environments via the term *g*_*ice*_ (4.2). *g*_*ice*_ is normally distributed 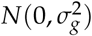, which corresponds to a compound symmetry model assuming a uniform covariance between genotypes in all environments. More sophisticated variance covariance (VCOV) structures are possible like the unstructured model using a specific covariance term for each pair of environments.

The error term *ϵ*_*ice*_ (4.3), is normally distributed *N*(0, ***R***) with within environment cross-specific variance error terms 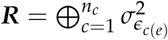. For an illustration purpose, the VCOV matrix of phenotypic observations coming from two different crosses in two different environments take the following form:

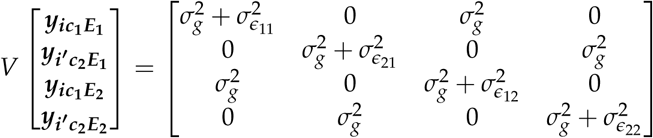

It represents a compound symmetry with heterogeneous cross-specific environment variances. In Model 4, the genotype by environment variance 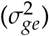 can not be distinguished from the error term 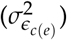. Model 4.4 is a matrix expression of model 4. The vector of phenotypic values ***y*** = [***y***_**1**_***y***_**2**_]^*T*^ includes the phenotypic observations of the two environments. The cross term ***X***_***c***_ ***β***_***c***_ and the QTL term ***X***_***Q***_ ***β***_***Q***_ are extended to model environment-specific cross and QTL effects.

### MPP-ME QTL detection methods

Here we present four methods to perform a QTL detection in MPP-ME experiments. The first three methods are two-stage analyses using first a phenotypic model to calculate genotype BLUEs that are used afterwards in a genotypic QTL detection model. The last method is a one-stage analysis. Method M1 performs a QTL analysis on genotype BLUEs calculated across environments. It uses the BLUEs calculated with model 2 as response variable in QTL model 3. Method M1 tests for association between the genotype and ***y***_***E*(1**,**2)**_, which represents a main phenotypic effect across environments. Therefore, M1 does not allow to estimate the QTLxE effects.

Method M2 allows to model the QTLxE interactions by performing separate QTL analyses in each environment. M2 uses the BLUEs calculated within each environment (***y***_***E*1**_, ***y***_***E*2**_) in QTL model 3. The methods M1 and M2 do not take the advantage of analyzing jointly the phenotype data measured in different environments, which has been shown to provide a greater understanding of the GxE interactions (Malosetti et al., 2004; Alimi et al., 2013). A proper MPP GxE QTL detection using mixed model allows to model the heterogeneity of genetic variance across environments and the genetic covariance between environments (Malosetti et al., 2013).

Method M3 analyzes jointly the within environment genotypes BLUEs (***y***_***E*1**_, ***y***_***E*2**_) using model 4 taking the covariance between the same genotype measured in different environments into consideration. The last method (M4) is a one-stage analysis on the plot data. For the one-stage analysis, we followed Boer et al. (2007) and separated the genetic effect terms of model 2 (*G*_*ic*_ + *GE*_*ice*_) into a check entries term (*G*_*i*_,_(*e*)_) and two terms to model the tested genotypes by the QTLs 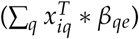 and the residual genetic effect (*g*_*ice*_). Method M4 allows to simultaneously estimates the non-genetic effects due to the experimental design (*des*_(*e*)_) and the QTL variations.

The variance of the error term *ϵ*_*ic*(*e*)*p*_ of models 1, 2 and M4 can be modelled by different VCOV structures taking for example spatial variations into consideration. In models 3, 4, and M4, the cross (*µ*_*ce*_) and the QTL terms (*β*_*qe*_) were fixed. The Wald test (McCulloch and Searle, 2001, 5.39) tests the global null hypothesis of all QTL allelic substitution effects being equal to zero. In models 4 and M4, the null hypothesis will be rejected if one allele is different from zero in at least one environment. The combination of four methods (M1-4) and three types of QTL effects (parental, ancestral, bi-allelic) represents 12 models for analyzing MPP-ME QTL experiments.

### Plant material

To illustrate our methodology, we focused on two examples showing significant and observable QTLxE interactions. The examples came from the Flint EU-NAM and the US-NAM populations tested for a single trait in two environments.

## EU-NAM data

The maize EU-NAM Flint population was composed of 811 double haploid (DH) lines coming from 11 crosses between UH007 and 11 peripheral parents representative of North Europe maize diversity (Bauer et al., 2013; Lehermeier et al., 2014). The EU-NAM Flint population was evaluated in six European locations for five traits. We used the raw phenotypic data provided by Lehermeier et al. (2014) http://www.genetics.org/content/198/1/3/suppl/DC1. We used the raw genotypic data provided by Bauer et al. (2013) available here http://www.ncbi.nlm.nih.gov/geo/query/acc.cgi?acc=GSE50558, and the consensus map from Giraud et al. (2014) available at: http://maizegdb.org/data_center/reference?id=9024747. After quality control, we kept 5949 markers spread on 10 chromosomes for a total map length of 1584 cM. For the ancestral model, we clustered the parental lines at each marker position using a two cM window around the marker with the R package clusthaplo (Leroux et al., 2014). We detected an average of 6.7 ancestral classes along the genome.

From the possible combinations of traits and environments, we focused on dry matter yield (DMY, decitons per hectare, 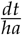) measured at La Coruña (Spain – CIAM) and at Roggenstein (Germany – TUM). Within a location, the trials were laid out as augmented p-rep designs with one third of the genotypes replicated. The genotypes were laid out with parents and checks in 160 incomplete block consisting of eight plots. Therefore, in model 1, 2 and M4 *des*(*e*) = *rep*_*l*(*e*)_ + *block*_*m*(*le*)_ to account for the *l*^*th*^ replicate, and the *m*^*th*^ block within replicate effects. The model 2 was equivalent to model 1 in Lehermeier et al. (2014). In M4, the variance of the error term was environment and cross specific (4.3). The average heritability on a line mean basis within cross over environments was 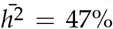 (supplementary material S1). The Pearson correlation between the two within environment BLUEs was equal to 0.35.

## US-NAM data

The maize US-NAM population was composed of 4421 recombinant inbred lines (RIL) coming from 25 crosses between B73 and 25 peripheral parents representative of the international maize diversity (McMullen et al., 2009). It was evaluated for 19 traits in up to 11 environments (Hung et al., 2012). We used the phenotypic data provided by Hung et al. (2012). The genotypic marker data and map data came from Ogut et al. (2015). We downloaded the data from http://www.panzea.org. We used the map with 1478 markers (1 mk/cM) and a total map length of 1434 cM. For the US-NAM data, we did not have the IBS marker data. Therefore, we did not calculate the ancestral and the bi-allelic models, and we restricted our analyses to the parental model.

We focused on days to anthesis (DTA) measured in 2007 in the North Carolina (NC) and the New York (NY) environments. Within an environment, the experimental design was a set design where each set contained all lines of a cross. Each set was randomized across environments as an *α*-design. The *α*-design was augmented by including the two parental lines of the cross within each incomplete block. The row and column effects were defined at the environment level (Hung et al., 2012). Therefore, in models 1, 2 and M4 *des*(*e*) = *set*_*l*(*e*)_ + *block*_*m*(*le*)_ + *row*_*n*(*e*)_ + *col*_*o*(*e*)_ to model the *l*^*th*^ set, *m*^*th*^ block within set, *n*^*th*^ row and *o*^*th*^ column effects. Model 2 was similar to the one used by Hung et al. (2012) removing the cross term. The set and the cross effects were not confounded because the parents and the checks were considered as being part of an ‘extra’ cross.

For the BLUEs computation in model 1 and 2, the VCOV structure of the error term was modelled by an environment-specific autoregressive correlation in the rows and columns (AR1 × AR1) (Gilmour et al., 1997). For the one-stage M4 analysis, the modelling of the spatial trend was however computationally too intensive, so we only used within environment cross-specific error terms (4.3). The average heritability on a line mean basis within cross over environments was 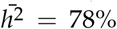 (supplemental material S1). The Pearson correlation between the two within environment BLUEs was equal to 0.25.

### QTL detection procedure

The QTL detection procedure was composed of a simple interval mapping scan to select cofactors followed by a composite interval mapping (CIM) scan to build a multi-QTL model. The final list of QTLs was evaluated using a backward elimination. The cofactors were selected with a minimum in between distance of 50 cM to avoid model overfitting. The QTLs were selected with a minimum distance of 20 cM. We fixed the cofactor and QTL detection thresholds to −*log*10(*p* − *value*) = 4. We performed the QTL detection extending the R package mppR (Garin et al., 2018) to the multi-environment situation. The mixed models were calculated using asreml-R (Butler et al., 2009). The R package programmed can be found here https://github.com/vincentgarin/mppGxE.

### Cross-validation

We adapted the cross-validation (CV) procedure described by Utz et al. (2000) to the MPP-ME context to evaluate the performances of the QTL detections. For each combination of methods (M1-4) and type of QTL effects (parental, ancestral, bi-allelic), we ran three times a three-fold CV procedure, which gave us nine estimates for each parameter. We did not perform the CV for the one-stage (M4) QTL analysis of the US-NAM data due to computational limitations. We partitioned the full dataset at the within-cross level into an estimation set (ES) and a test set (TS). We used the estimated effects of the QTLs detected in the ES to predict phenotypic values of the ES and TS (e.g. 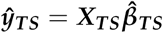).

We calculated the proportion of genetic variance explained in the ES by the 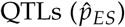 using the Pearson correlation between the reference values ***y***_***ES***_ and the predicted values ***ŷ***_***ES***_. We calculated the proportion of genetic variance predicted in the TS by the 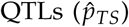 using the Pearson correlation between the reference values ***y***_***TS***_ and the predicted values ***ŷ***_***TS***_. We used as reference values (***y***_***ES***_ and ***y***_***TS***_) the within environment BLUEs (model 1) to compare our results across methods. The 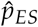 and 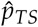 were computed within crosses per environment. We estimated the 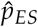 and 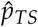 at the whole MPP level by calculating the average within cross values.

### Modelling QTL effect in relation to environmental information

The methods M2, M3 and M4 allow to detect environmental differences of the QTL effects but they do not allow to understand how environmental characteristics influence the QTL effects. A natural extension is to integrate environmental information to understand better the QTLxE interaction. Inspired by the model 16 proposed by Malosetti et al. (2013), we can reformulate the QTL part of *G*_*ijk*_ (4.1) to describe the QTL effect of a single QTL (*q*\*) in term of an environmental covariate (*Z*_*e*_).

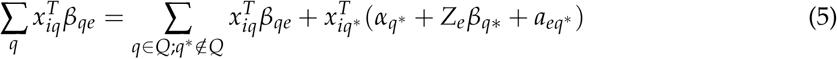

In this formula the QTL part is the same as in 4.1 for all QTLs except for *q*\*. For the QTL *q*\*, we decompose its environmental effect (*β*_*q*\**e*_) into a main effect component *α*_*q*\*_ and a component *β*_*q*\*_ describing the sensitivity to the environmental covariate *Z*_*e*_. In the MPP context, *α*_*q*\*_ and *β*_*q*\*_ will be vectors of dimension [1 × *n*_*al*_] containing one element per QTL allele *l. α*_*q*l*_ represents the QTL effect of allele *l* when *Z*_*e*_ is equal to zero. *β*_*q*\**l*_ is the environmental sensitivity of QTL allele *l* and represents the amount of change in trait quantity for one extra unit of *Z*_*e*_. *α*_*q*l*_ and *β*_*q*\**l*_ are both defined with respect to a reference allele (e.g. the central parent). *Z*_*e*_ is the value of the environmental covariate (e.g. temperature). Finally, 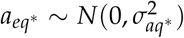 is the residual unexplained QTL effect.

## III. RESULTS

### Cross-validation results

Table 2 contains the CV results for the two populations over the different combinations of methods (M1-M4) and QTL effects (parental, ancestral, bi-allelic). We did not detect large differences in terms of prediction power 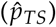 between the different methods. For example, for the US-NAM data in the first environment (NC), the average 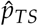 across crosses was equal to 15, 16 and 17 for the M1, M2 and M3 methods, respectively. Similarly, for the EU-NAM ancestral model in the second environment (TUM), the 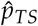 was equal to 11, 11, 15 and 12 for the M1 to M4 methods, respectively. Concerning the number of QTLs, we observed that, on average, the separate single environment M2 method detected the lowest number of QTLs. The one-stage M4 analyses detected less QTLs than M1 and M3. In some cases, method M1 detected the largest number of QTLs like in the three EU-NAM complete data analyses.

**Table 2:** Cross-validation results of the EU-NAM and US-NAM data for each combination of methods (M1-4) and type of QTL effects (parental, ancestral bi-allelic). Number of QTLs detected in the complete data analyses (N.QTL). Average number of detected QTLs in the TS over the CV runs (N.QTLcv). Average proportion of variance explained by the detected QTLs in the estimation set 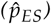 over the crosses with minimum and maximum within cross values. Average proportion of variance predicted by the detected QTLs in the test set 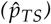 over the crosses with minimum and maximum within cross values.

|  |  | EU-NAM |  |  |  | US-NAM |  |  |  |
| --- | --- | --- | --- | --- | --- | --- | --- | --- | --- |
| | | $N.QTL$ | $N.QTL_{cv}$ | $\hat{p}_{ES}$ | $\hat{p}_{TS}$ | $N.QTL$ | $N.QTL_{cv}$ | $\hat{p}_{ES}$ | $\hat{p}_{TS}$ |
| parental |  |  |  |  |  |  |  |  |  |
| M1 | E1 | 8 | 4.9 | 23 (13-42) | 10 (5-19) | 22 | 14.8 | 29 (15-45) | 15 (4-37) |
| M1 | E2 | 8 | 4.9 | 21 (6-44) | 9 (2-19) | 22 | 14.8 | 18 (4-47) | 9 (1-35) |
| M2 | E1 | 6 | 3.7 | 23 (13-53) | 8 (2-36) | 24 | 16.6 | 40 (27-56) | 16 (4-31) |
| M2 | E2 | 3 | 2.7 | 20 (6-35) | 12 (3-25) | 7 | 4.8 | 14 (3-45) | 8 (1-39) |
| M3 | E1 | 7 | 5.1 | 28 (15-54) | 10 (2-24) | 27 | 17.2 | 40 (27-55) | 17 (6-38) |
| M3 | E2 | 7 | 5.1 | 33 (11-61) | 12 (3-21) | 27 | 17.2 | 27 (15-54) | 8 (0-33) |
| M4 | E1 | 6 | 3.7 | 21 (10-39) | 12 (3-24) | 18 |  |  |  |
| M4 | E2 | 6 | 3.7 | 22 (7-34) | 16 (1-28) | 18 |  |  |  |
| ancestral |  |  |  |  |  |  |  |  |  |
| M1 | E1 | 9 | 5.2 | 17 (4-36) | 11 (1-24) |  |  |  |  |
| M1 | E2 | 9 | 5.2 | 19 (7-33) | 11 (3-29) |  |  |  |  |
| M2 | E1 | 6 | 4.6 | 21 (8-58) | 10 (3-20) |  |  |  |  |
| M2 | E2 | 4 | 3.8 | 21 (8-41) | 11 (1-19) |  |  |  |  |
| M3 | E1 | 6 | 7.2 | 28 (6-59) | 10 (1-23) |  |  |  |  |
| M3 | E2 | 6 | 7.2 | 30 (6-57) | 15 (4-30) |  |  |  |  |
| M4 | E1 | 7 | 3.7 | 15 (5-24) | 10 (3-17) |  |  |  |  |
| M4 | E2 | 7 | 3.7 | 19 (4-33) | 12 (3-24) |  |  |  |  |
| bi-allelic |  |  |  |  |  |  |  |  |  |
| M1 | E1 | 12 | 8.4 | 18 (5-28) | 11 (2-31) |  |  |  |  |
| M1 | E2 | 12 | 8.4 | 20 (7-32) | 12 (6-30) |  |  |  |  |
| M2 | E1 | 7 | 5.7 | 20 (7-32) | 10 (3-21) |  |  |  |  |
| M2 | E2 | 5 | 5.1 | 20 (7-33) | 10 (1-22) |  |  |  |  |
| M3 | E1 | 10 | 7.8 | 20 (6-34) | 9 (1-14) |  |  |  |  |
| M3 | E2 | 10 | 7.8 | 19 (3-35) | 14 (7-23) |  |  |  |  |
| M4 | E1 | 9 | 7 | 15 (4-27) | 12 (2-25) |  |  |  |  |
| M4 | E2 | 9 | 7 | 18 (3-32) | 12 (3-25) |  |  |  |  |

Concerning the type of QTL effect (parental, ancestral, bi-allelic), we also did not detect any difference in terms of prediction power 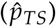. For the EU-NAM data, the average within crosses 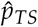 of the parental, of the ancestral, and of the bi-allelic model varied between: 8-16, 10-15, and 9-14, respectively. We noticed however, that we detected on average more QTLs with the bi-allelic and the ancestral models.

### −log10(p-values) scatter-plots

In Figure 1, we plotted the −log10(p-values) of the CIM profiles obtained with the method M4 with respect to the methods M1 to M3 for all complete data QTL analyses. We could observe that, in general, the −log10(p-values) were larger in M4. The differences between the M4 and the M2 profiles were the largest. Concerning M4 versus M1, an important fraction of the −log10(p-values) were superior in the M4 profiles with respect to the M1 profiles. However, for the most significant −log10(p-values), the M1 method gave sometimes slightly larger −log10(p-values) than the M4 method, for example for the EU-NAM parental model. The −log10(p-values) obtained with M3 and M4 were similar.

**Figure 1:**
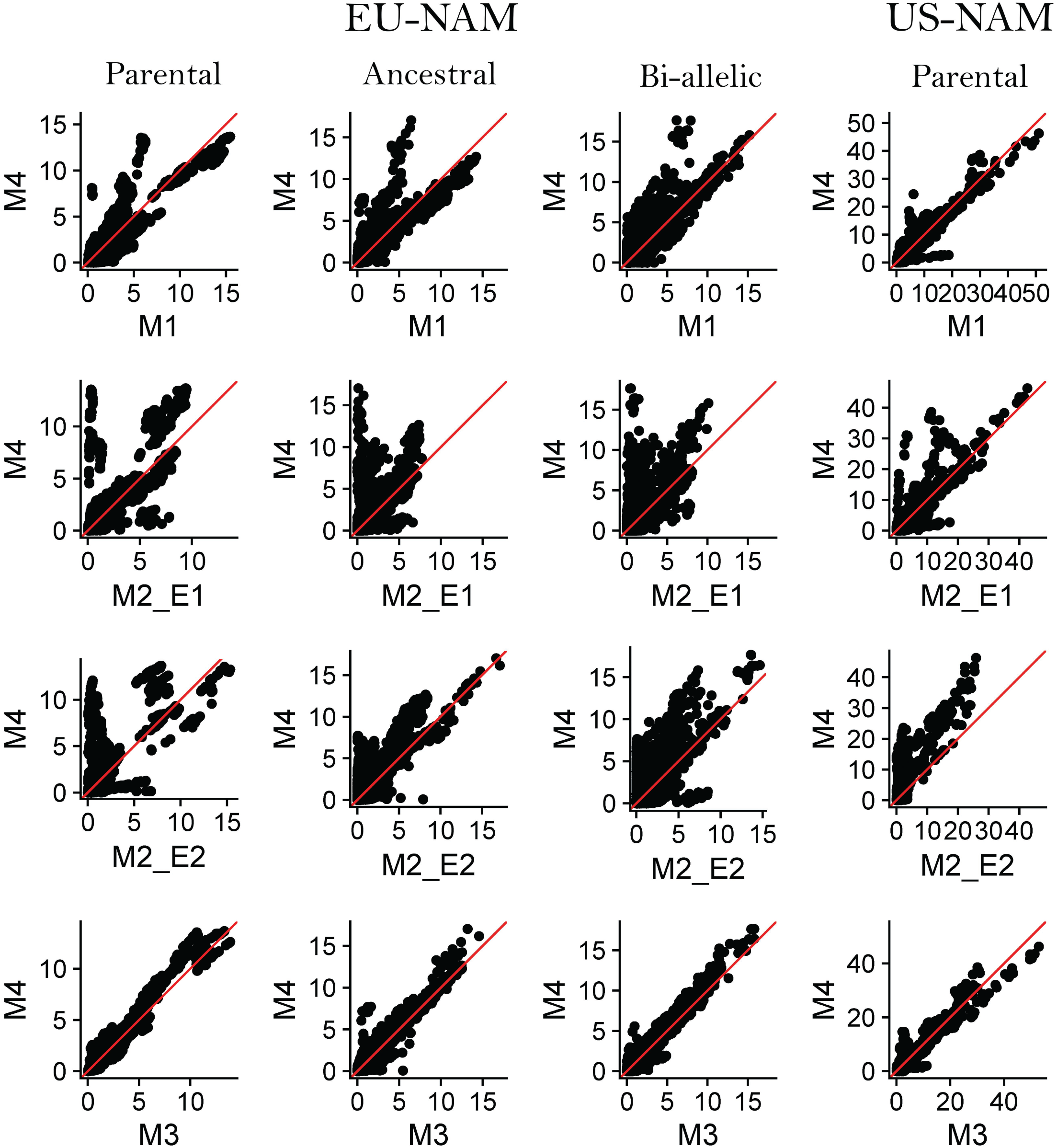
CIM −log10(p-values) scatter-plots of methods M1-M3 compared to M4 for both the EU-NAM and the US-NAM whole population QTL analyses.

### Detected QTLs

In Figure 2, we plotted the −log10(p-values) CIM profile of the US-NAM M4 parental QTL analysis with a representation of the genetic effect significance per parent and per environment along the genome. On that plot, we observed that the QTLs on chromosomes eight, nine, and ten were detected with a large significance. Looking at the genetic effect distribution along the genome, we also noticed that these QTLs potentially had a different parental allelic series between the environments. The information about the significance of the QTL genetic effect along the genome should however be taken with caution because it is based on an incremental and conditional Wald test that can change given the order of the tested parameters (Butler et al., 2009).

**Figure 2:**
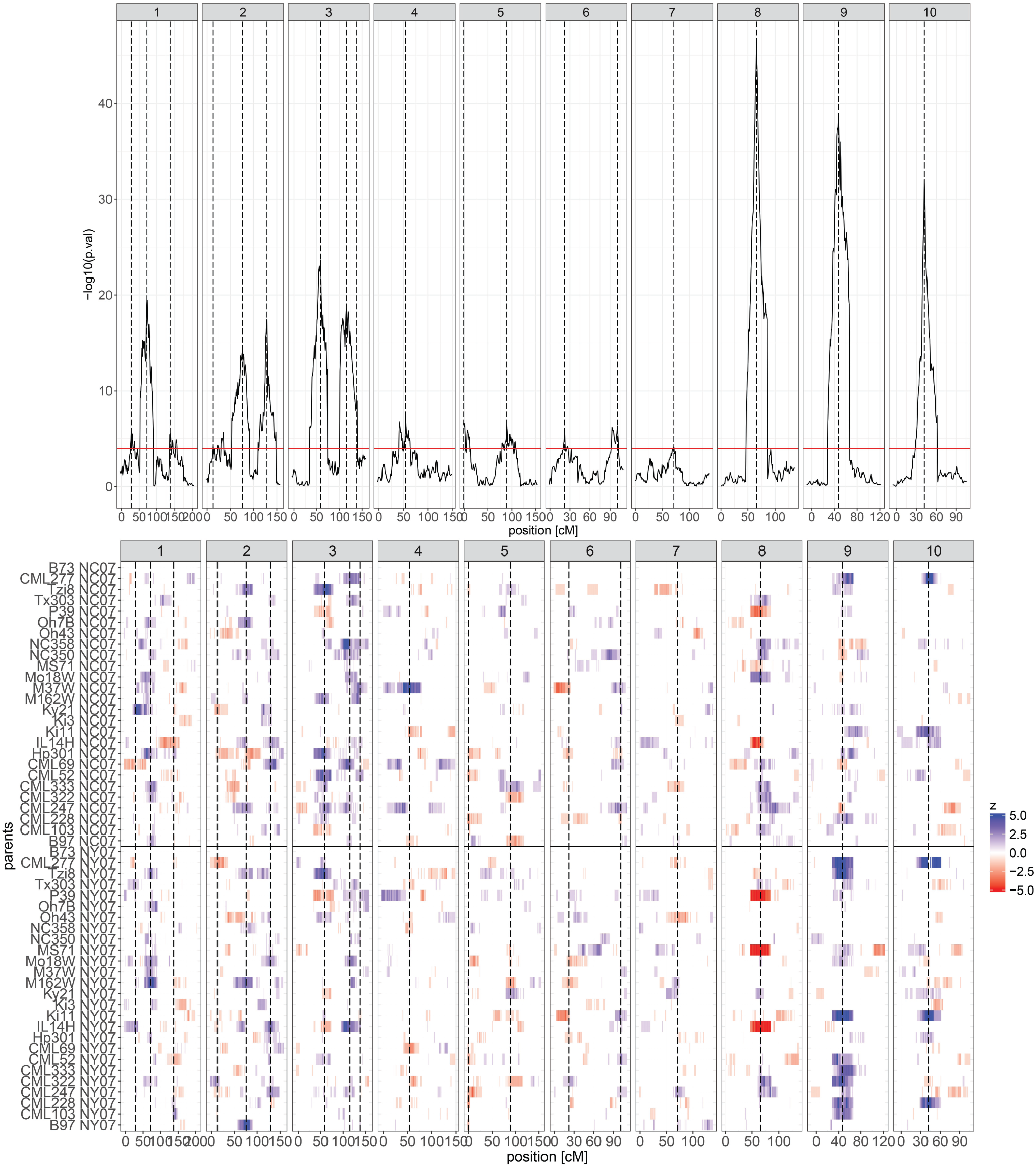
CIM −log10(p-values) profile of the US-NAM M4 parental QTL analysis. The lower part of the figure represents the within environment parental QTL allelic significance along the genome. The Wald test p-values of the parental allelic substitution effects are converted into a colour code from > 0.05 (1) to > 10^−5^ (5). The colours red (negative) and blue (positive) correspond to the sign of the QTL effect

In Figure 3, we plotted the −log10(p-values) CIM profile of the EU-NAM M4 ancestral QTL analysis with a representation of the genetic effect significance per parent and per environment along the genome. We noticed that the QTL on chromosome six had an interesting allelic series. Indeed, many parents were grouped in the same ancestral class and the QTL had a genetic effect specific to the second environment (TUM). The rest of the detected positions for all methods and type of QTL model combinations can be found in the supplementary material S2.

**Figure 3:**
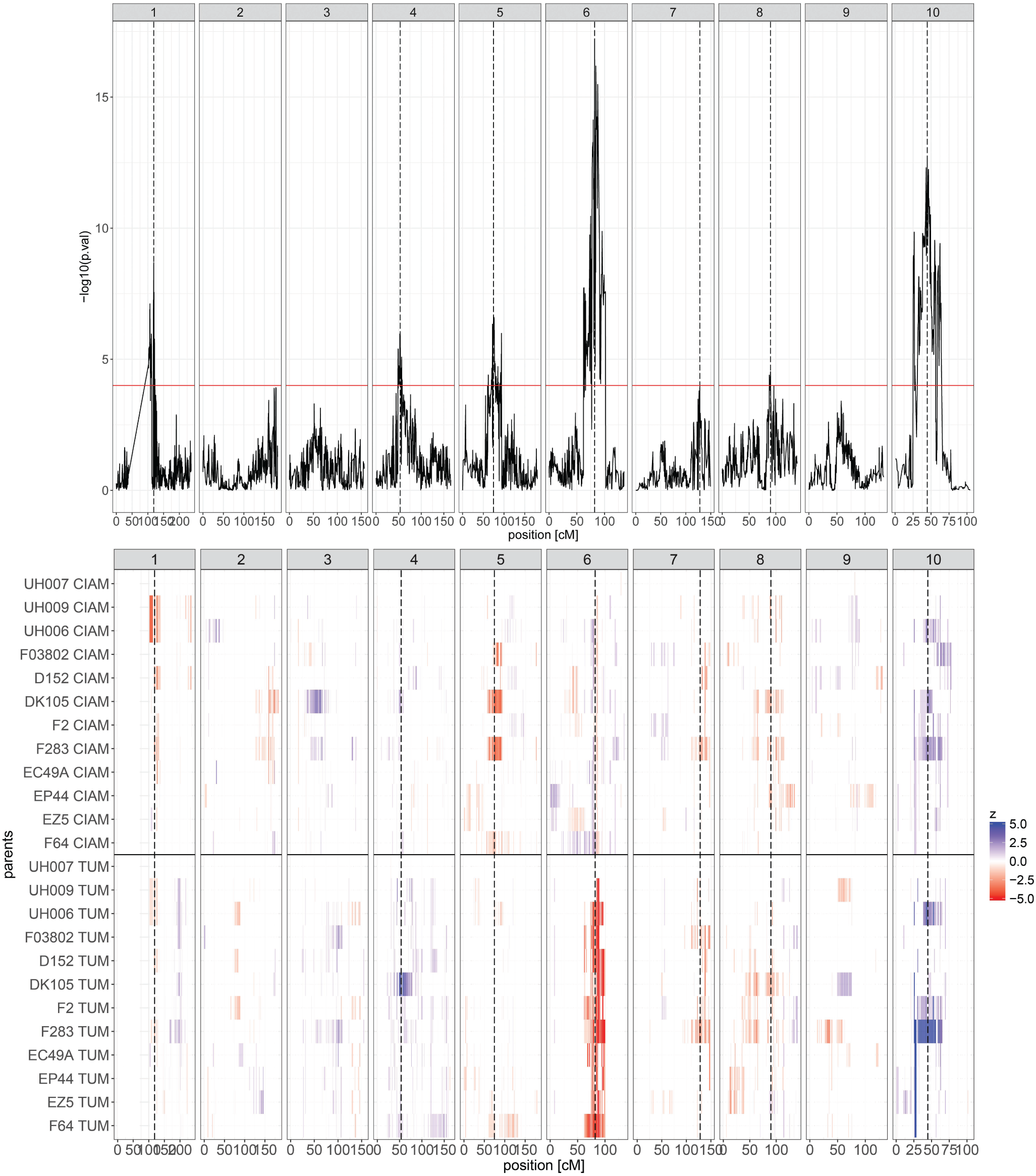
CIM −log10(p-values) profile of the EU-NAM M4 ancestral QTL analysis. The lower part of the figure represents the within environment parental QTL allelic significance along the genome. The Wald test p-values of the parental allelic substitution effects are converted into a colour code from > 0.05 (1) to > 10^−5^ (5). The colours red (negative) and blue (positive) correspond to the sign of the QTL effect

### Estimation of the QTL allelic substitution effects

We represented the QTL allelic series of the QTL detected on chromosome six in the EU-NAM ancestral model and the QTLs detected on chromosomes eight, nine and ten in the US-NAM data in Figure 4. The estimated QTL effects were conditioned on the list of cofactors detected with the corresponding final model. The numerical QTL allelic substitution effect values can be found in the supplementary material S3. In the text, the standard errors of the allelic effects are given in parentheses. In Figure 4, we observed the differences in terms of QTL effect estimation between method M1 using BLUEs representing an average phenotypic effect across environments, and the method M4, which allows to estimate environment-specific QTL effects. The results obtained with method M4 were comparable to the ones obtained with methods M2 and M3. A full comparison between the estimated QTLs effects obtained with all the methods can be found in the supplementary material S4.

**Figure 4:**
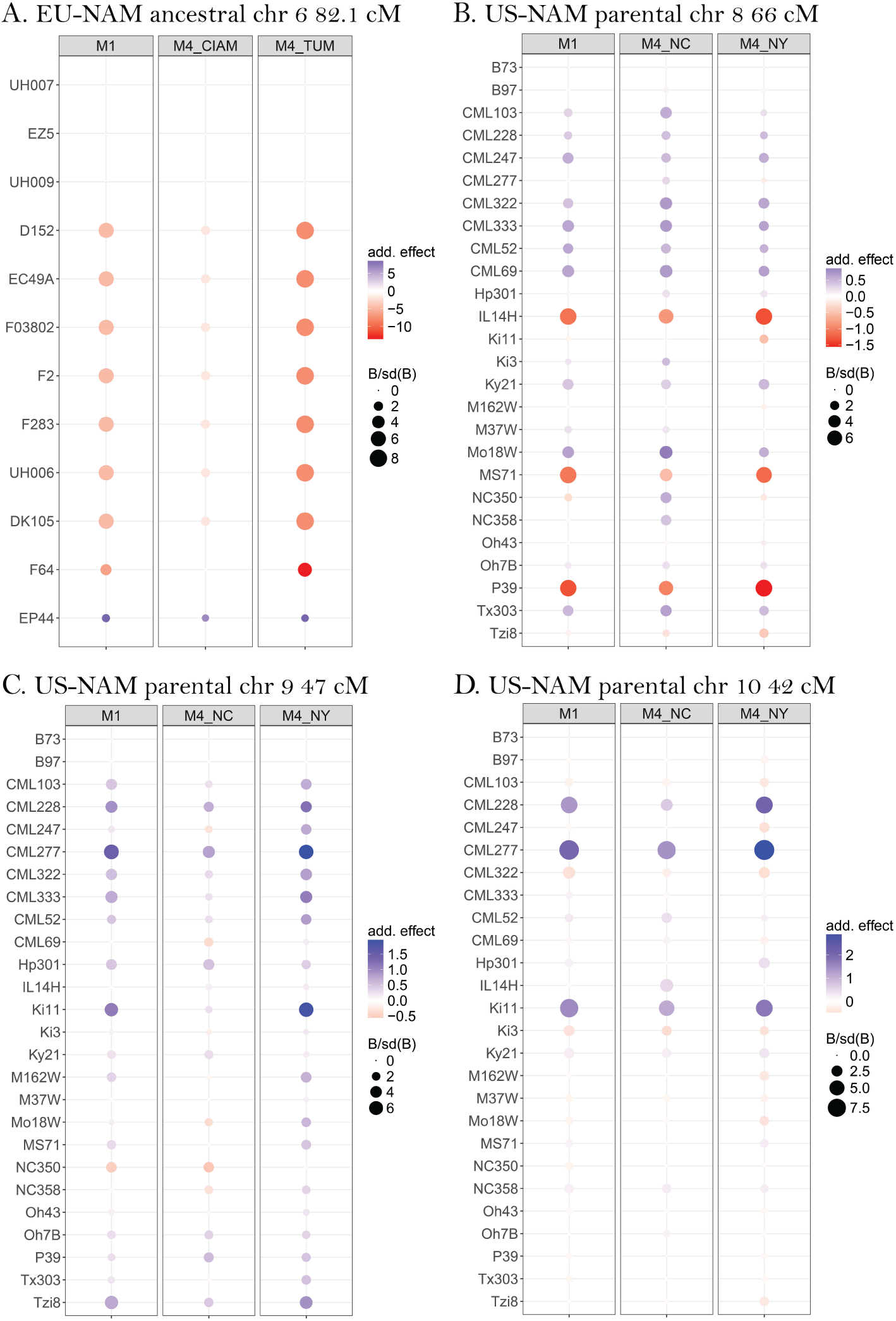
Comparison of the allelic substitution effect series between M1 and M4 for four QTL positions: A) EU-NAM ancestral model chromosome 6 82.1 cM; B) US-NAM parental model chromosome 8 66 cM; C) US-NAM parental model chromosome 9 47 cM; D) US-NAM parental model chromosome 10 42 cM. The colour intensities are proportional to the allelic effect. The allelic effects are deviations in decitons per hectare (A) and days (B-D) with respect to the central parent (UH007 or B73). The sizes of the dots are proportional to the ratio between the allelic effect and its standard error.

The QTL detected on chromosome six with the ancestral model in the EU-NAM population (Figure 4-A), was the most illustrative example of environment-specific QTL effects. In the second environment (TUM), an ancestral allele inherited by parents D152, EC49A, F03802, F2, F283, UH006 and DK105 had a strong negative effect of 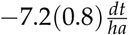 with respect to the ancestral group containing parents UH007, EZ5, and UH009. In the first environment (CIAM), the effect of the main ancestral group was substantially reduced 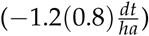 and non-significant. In the M1 method, the main ancestral allele took an average values 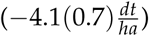 across the two environments.

In example 4B, we observed that the three most significant parental alleles had an effect that was stronger in the second environment (NY) compared to the first one (NC). The estimated allelic substitution effects (in days) of parental alleles IL14H, MS71, and P39 were −1.3(0.1) vs −0.8(0.1), −1.2(0.1) vs −0.5(0.1), and −1.6(0.2) vs −1.0(0.2), respectively. The estimated QTL effects obtained with M1 were again approximately averaged between the two environments with −1.1, −1.1 and −1.3 days respectively. Similarly, in example 4C, we observed that several parental alleles had a stronger positive expression in NY compared to NC. For example, the allelic substitution effect of Ki11 was equal to +1.9(0.2) days in NY and +0.3(0.2) days in NC. The allelic substitution effect of CML277 was equal to +1.9(0.2) days in NY while it was equal to +0.8(0.2) days in NC. Finally, in example 4D, the most significant parental alleles had a stronger effect in the second (NY) environment. For example, the estimated allelic substitution effect of CML277 was equal to +3(0.2) days in NY and +1.4(0.2) days in NC. Once again, in the examples 4C and 4D, the allelic substitution effects estimated with method M1 were approximately averaged across the two environments.

### QTL effect in relation to environmental information

To illustrate the extension of our models with environmental covariates, we re-analyzed the effect of the QTL detected on chromosome six (84.2 cM) in the EU-NAM with the M3 ancestral model including the effect of water precipitation (*Z*_*e*_). This QTL had five alleles: allele A (UH007-central parent), allele B (D152, EC49A, EP44, F2, F64, UH006), allele C (F03802, F283), allele D (UH009, DK105), and allele E (EZ5). We used the final QTL model and the average water precipitation in mm (*Wat*.(*mm*)) at each location between July and August obtained from https://en.climate-data.org/. To increase the range of the environmental covariate, we included four environments (La Coruña, Roggenstein, Einbeck, Ploudaniel). We considered the precipitation of the driest location (La Coruña – 35 mm) as the reference level. The precipitation in the other environments were expressed as the difference with respect to the reference.

In Table 3, we can observe the estimates of the QTL main effects (*α*) and QTL × environment effects (*β*) on the trait. We noticed that, in the driest environment (La Coruña), the allele B reduced the yield by 0.75 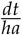 compared to allele A. When the level of precipitation increased, this difference was accentuated by 0.06 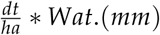. In Table 4, we calculated the difference in yield between an homozygous genotype with allele A versus B in the four environments (2 *(*α*_*B*_ + *Z*_*e*_ * *β*_*B*_)). In this table, we observed that the difference was equal to 1.5 in the driest reference environment (La Coruña). The extra yield given by allele A increased with more precipitation (e.g. 6.6 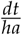 at Roggenstein).

**Table 3:** Parameter estimates of the environment-specific QTL effects expressed in terms of the QTL sensitivity to average water precipitation during July and August.

|  | Estimates | Std. Err | Units | Wald | Df | P(Wald) |  |
| --- | --- | --- | --- | --- | --- | --- | --- |
| $\alpha_A$ | 0 | | | 0 | 0 | - | |
| $\alpha_B$ | -0.75 | 1.07 | | 12.5 | 1 | <0.001 | *** |
| $\alpha_C$ | 2.02 | 1.39 | $dt * ha^{-1}$ | 3.5 | 1 | 0.06 | . |
| $\alpha_D$ | 1.82 | 1.44 | | 1.1 | 1 | 0.3 | |
| $\alpha_E$ | -9.02 | 4.6 | | 2.7 | 1 | 0.1 | |
| $\beta_A$ | 0 | | | 0 | 0 | - | |
| $\beta_B$ | -0.06 | 0.03 | | 4 | 1 | 0.05 | * |
| $\beta_C$ | -0.12 | 0.04 | $dt * ha^{-1} * Wat.(mm)$ | 8.4 | 1 | 0.003 | ** |
| $\beta_D$ | -0.03 | 0.04 | | 0.7 | 1 | 0.42 | |
| $\beta_E$ | 0.17 | 0.14 | | 1.6 | 1 | 0.21 | |

**Table 4:** Extra yield homozygous genotype with allele A versus B given water precipitation.

| Location | $Z_e$ (mm) | Yield (Std. Err) [$dt * ha^{-1}$] |
| --- | --- | --- |
| La Coruna | 0 | 1.50 (2.13) |
| Roggenstein | 42 | 6.60 (1.64) |
| Ploudaniel | 28 | 4.80 (1.35) |
| Einbeck | 33 | 5.46 (1.40) |

## IV. DISCUSSION

Several MPPs have been characterized in multiple environments but, most of the time, the QTL analyses were performed on genotype BLUEs calculated across environments, which represent average phenotypic values (Giraud et al., 2014; Buckler et al., 2009). We called that method M1. M1 does not estimate the QTLxE effects. Therefore, M1 does not allow to use the full information potential of MPP-ME QTL experiments, measuring QTL variations: a) within the MPP between sub-populations due to different genetic backgrounds; and b) between environments.

An alternative to estimate the QTLxE effects in MPPs is to perform separate QTL analyses within each environment (M2). However, this method does not model the covariance due to the repeated measurements on the same genotype. Therefore, we proposed two methods (M3 and M4) that allowed to model properly the QTLxE effects in MPPs. Those methods used simultaneously the phenotypic data from multiple environments taking into consideration the covariance existing between the same genotype measured in different environments. With respect to M3, M4 also integrated the sources of variation due to experimental design elements performing a one-stage analysis on the plot data. Using M3 and M4 we were able to detect important QTLs and we showed that these QTLs had environmental specific allelic effects. Finally, we extended our models to integrate environmental information and better characterize the QTLxE effects.

### Comparison of the detected QTLs with the existing literature

Using the different methods presented, we detected several interesting QTLs for DMY and for DTA. The QTL on chromosome six at 82.1 cM detected by the ancestral model in the EU-NAM population (Figure 4A) was also detected by Giraud et al. (2014). They detected a QTL at 83.5 cM, 90.5 cM and 90.7 cM using the connected, the 2cM-LDLA and the 1mk-LDLA models, respectively.

The QTLs detected for DTA in the US-NAM population on chromosomes eight, nine, and ten, at 66, 47, and 42 cM, respectively (Figure 4B, C and D), were also detected by Buckler et al. (2009). They detected corresponding QTLs on chromosomes eight, nine, and ten, at 67, 45.2, and 42.9 cM, respectively. According to Buckler et al. (2009), the QTL on chromosome eight is close to the vegetative to generative transition 1 (vgt1) gene (Salvi et al., 2002). Concerning the QTL on chromosome ten (42 cM), Giraud et al. (2014) also detected a QTL with a strong effect on flowering time between 45 to 50 cM. According to them, it corresponds to the ZmCCT gene Ducrocq et al. (2009).

### Estimation of the QTLxE effects

In Figure 4 and appendix S4, we showed that important QTLs had environment-specific allelic substitution effects. In those cases, the QTL effects estimated with method M1 were inaccurate. Those QTL effects were overestimated in one environment and underestimated in the other. Therefore, in presence of QTLxE effect, the use of method M2, M3, or M4 is necessary to estimate properly the QTL environmental differences.

The QTL detected on chromosome six at 82.1 cM in the EU-NAM population (Figure 4A) is a good illustration of the possibility to estimate QTL effect variations at two levels: within the MPP between sub-populations, and between environments. At that position, the QTL effect was rather consistent within the MPP because an important group of parents showed a negative effect that could be due to a common ancestral allele. For that QTL, we could also observe environmental variation because the allelic effects were only significant in the second environment. This example illustrates the interest of using a sound MPP GxE QTL detection methodology.

### QTL effect in relation to environmental information

In Tables 3 and 4, we showed that methods M3 or M4 could be extended to integrate environmental information to better characterize the QTLxE effects. The estimation of the water precipitation effect on a single QTL was a simple case with a unique environmental covariate but we could imagine more complex models with more QTLs and/or environmental covariates (Malosetti et al., 2004). We assumed a linear relationship between the QTL effect and the environmental covariate. More complex relationships such as a quadratic form or splines could also be assumed (van Eeuwijk et al., 2007). The ultimate goal of such an approach is to unravel the physiological mechanisms behind the QTL effects. This possibility to integrate environment information make methods M3 or M4 more attractive than M2.

### Full data analyses and cross-validation results

In terms of prediction power, the CV results (Table 2) did not allow to make a difference between the four methods. The 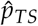 were mostly similar. We showed in the full data analyses and the CV that, on average, method M2 detected less QTLs. The prediction power 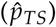 of M2 was also reduced with respect to the other methods, even if the differences were small. According to our results, the use of separate within environment analyses is therefore not an optimal strategy. The joint analysis of multi-environment data, as performed in M3 and M4, accounted better for the shared effects across environments, which was beneficial for the QTL analysis.

In the full data analysis, we also noticed that in all EU-NAM analyses (parental, ancestral, bi-allelic), method M1 detected the largest number of QTLs. In those cases, the extra power of M1 could be explained by the fact that this method uses a reduced number of degrees of freedom (df) to estimate the QTL effect. Indeed, in M1, the QTL term uses *n*_*al*_ − 1 df, while in M2, M3 or M4, the QTL term uses *N*_*Env*_ * (*n*_*al*_ −1) dfs. When the QTLxE effect is strong, the loss in power due to the extra df for the QTLxE term is compensated by a better modelling of the GxE effect. However, when the QTL effect is consistent across the environments, the additional modelling of the QTLxE variation penalizes the test for a QTL. With consistent QTL effects, method M1 is more parsimonious.

We would like to emphasize that we selected examples with significant observable GxE interactions but that these situations did not represent the majority of the cases we tested. This observation is in agreement with Buckler et al. (2009) who found that, for flowering time traits in the US-NAM population, the environment-specific QTL effects were small compared to the main QTL effects across environments. The potential weakness of QTLxE effects with respect to the main QTL effects in the US-NAM and EU-NAM populations, could explain why, in some cases, method M1 obtained better results than the GxE analyses (M2-M4). The existence of GxE effects could be more important in MPP-ME QTL experiments where the environmental conditions are more contrasted, for example, in experiments testing the same population in control versus heat or salt stress conditions (Saade et al., 2016).

As observed in Table 2, method M3 detected on average more QTLs than M4. Looking at the list of detected QTLs (appendix S2), we noticed that, in the EU-NAM analyses, the positions of the QTLs in M3 and M4 were consistent. In the US-NAM analyses, the positions of the QTLs with a large significance were consistent between M3 and M4. However, several QTLs with a low or medium significance were either only detected in M3, or distant by 10-15 cM between the two methods.

The difference between M3 and M4 could be explained by the modelling of the experimental design variation in M4. Piepho and Pillen (2004) showed that even if the variance of the experimental design factors were smaller than other elements like the genetic covariance, it could still reduce substantially the QTL effect when it was used in a one-stage analysis. This reduction of the QTL effect could explain why we detected less QTLs in M4 with respect to M3. We should still remember that in the US-NAM analysis we used an AR1 × AR1 covariance structure for the error term to calculate the within environment BLUEs used in the M3 analyses. Due to computational limitations, we did not use the AR1 × AR1 covariance structure in M4. This could also explain the differences between M3 and M4 in the US-NAM analyses.

### Other extensions

To present our methodology, we used examples from NAM populations but our methods and the reasoning behind it are also valid for any MPP composed of crosses like diallel population or factorial designs used in breeding programs. Our methods can also be adapted to the multi-trait situation. The analysis of longitudinal traits measured at different time points using a VCOV reflecting the time dependence could be a possibility. For an illustration purpose, we only used data coming from two environments but we could increase this number. However, the fitting of mixed models on large datasets can be computationally intensive. For example, it took us four days to perform the M4 one-stage QTL detection in the US-NAM population on a personal computer (Intel Core i7-3770 CPU 3.4 GHz).

## Conclusions

We proposed mixed model methods to detect QTL in MPP-ME data. These models analyzed jointly data from several environments and modelled the genetic covariance due to repeated measurements on the same genotype. Our methods allowed to estimate the QTLxE effect while the methods using genotype BLUEs calculated across environments did not. However, the methods focusing on the main QTL effects remain useful if the QTL effects are consistent across the environments. Moreover, we showed that our methods could be extended to integrate environmental information and understand better the mechanisms behind the QTLxE effects. The methods we proposed are therefore an interesting tool to exploit the full information potential of MPP-ME data. They allow to estimate the QTL variations: a) within MPP between sub-populations due to different genetic backgrounds; and b) between environments.

## Notes

https://github.com/vincentgarin/mppGxE_data

## REFERENCES

Alimi, N., Bink, M., Dieleman, J., Magán, J., Wubs, A., Palloix, A., and van Eeuwijk, F. A. (2013). Multi-trait and multi-environment QTL analyses of yield and a set of physiological traits in pepper. Theor. Appl. Genet., 126(10):2597–2625.

Bauer, E., Falque, M., Walter, H., Bauland, C., Camisan, C., Campo, L., Meyer, N., Ranc, N., Rincent, R., Schipprack, W., et al. (2013). Intraspecific variation of recombination rate in maize. Genome Biol., 14(9):R103.

Boer, M. P., Wright, D., Feng, L., Podlich, D. W., Luo, L., Cooper, M., and van Eeuwijk, F. A. (2007). A mixed-model quantitative trait loci (QTL) analysis for multiple-environment trial data using environmental covariables for qtl-by-environment interactions, with an example in maize. Genetics, 177:1801–1813.

Broman, K. W., Wu, H., Sen, Ś., and Churchill, G. A. (2003). R/qtl: Qtl mapping in experimental crosses. Bioinformatics, 19(7):889–890.

Buckler, E. S., Holland, J. B., Bradbury, P. J., Acharya, C. B., Brown, P. J., Browne, C., Ersoz, E., Flint-Garcia, S., Garcia, A., Glaubitz, J. C., et al. (2009). The genetic architecture of maize flowering time. Science, 325(5941):714–718.

Butler, D., Cullis, B. R., Gilmour, A., and Gogel, B. (2009). Asreml-r reference manual. The State of Queensland, Department of Primary Industries and Fisheries, Brisbane.

Cavanagh, C., Morell, M., Mackay, I., and Powell, W. (2008). From mutations to MAGIC: resources for gene discovery, validation and delivery in crop plants. Curr. Opin. Plant Biol., 11(2):215–221.

Ducrocq, S., Giauffret, C., Madur, D., Combes, V., Dumas, F., Jouanne, S., Coubriche, D., Jamin, P., Moreau, L., and Charcosset, A. (2009). Fine mapping and haplotype structure analysis of a major flowering time quantitative trait locus on maize chromosome 10. Genetics, 183(4):1555–1563.

Garin, V., Wimmer, V., Borchardt, D., van Eeuwijk, F. A., and Malosetti, M. (2018). mppR: Multi-Parent Population QTL Analysis. R package version 1.1.10.

Garin, V., Wimmer, V., Mezmouk, S., Malosetti, M., and van Eeuwijk, F. A. (2017). How do the type of QTL effect and the form of the residual term influence QTL detection in multi-parent populations? a case study in the maize EU-NAM population. Theor. Appl. Genet., 130(8):1753–1764.

Gilmour, A. R., Cullis, B. R., and Verbyla, A. P. (1997). Accounting for natural and extraneous variation in the analysis of field experiments. J Agric Biol Environ Stat, 2(3):269–293.

Giraud, H., Lehermeier, C., Bauer, E., Falque, M., Segura, V., Bauland, C., Camisan, C., Campo, L., Meyer, N., Ranc, N., et al. (2014). Linkage disequilibrium with linkage analysis of multiline crosses reveals different multiallelic qtl for hybrid performance in the flint and dent heterotic groups of maize. Genetics, 198(4):1717–1734.

Hung, H., Browne, C., Guill, K., Coles, N., Eller, M., Garcia, A., Lepak, N., Melia-Hancock, S., Oropeza-Rosas, M., Salvo, S., et al. (2012). The relationship between parental genetic or phenotypic divergence and progeny variation in the maize nested association mapping population. Heredity, 108(5):490–499.

Jourjon, M.-F., Jasson, S., Marcel, J., Ngom, B., and Mangin, B. (2005). Mcqtl: multi-allelic QTL mapping in multi-cross design. Bioinformatics, 21(1):128–130.

Korte, A., Vilhjálmsson, B. J., Segura, V., Platt, A., Long, Q., and Nordborg, M. (2012). A mixed-model approach for genome-wide association studies of correlated traits in structured populations. Nat. Genet., 44(9):1066–1071.

Lehermeier, C., Krämer, N., Bauer, E., Bauland, C., Camisan, C., Campo, L., Flament, P., Melchinger, A. E., Menz, M., Meyer, N., et al. (2014). Usefulness of multiparental populations of maize (Zea *mays* L.) for genome-based prediction. Genetics, 198(1):3–16.

Leroux, D., Rahmani, A., Jasson, S., Ventelon, M., Louis, F., Moreau, L., and Mangin, B. (2014). Clusthaplo: a plug-in for mcqtl to enhance QTL detection using ancestral alleles in multi-cross design. Theor. Appl. Genet., 127(4):921–933.

Malosetti, M., Ribaut, J.-M., and van Eeuwijk, F. A. (2013). The statistical analysis of multi-environment data: modeling genotype-by-environment interaction and its genetic basis. Front. Physiol., 4:44.

Malosetti, M., Voltas, J., Romagosa, I., Ullrich, S., and van Eeuwijk, F. A. (2004). Mixed models including environmental covariables for studying qtl by environment interaction. Euphytica, 137(1):139–145.

McCulloch, C. E. and Searle, S. R. (2001). Generalized, linear, and mixed models. Wiley Online Library.

McMullen, M. D., Kresovich, S., Villeda, H. S., Bradbury, P., Li, H., Sun, Q., Flint-Garcia, S., Thornsberry, J., Acharya, C., Bottoms, C., et al. (2009). Genetic properties of the maize nested association mapping population. Science, 325(5941):737–740.

Ogut, F., Bian, Y., Bradbury, P. J., and Holland, J. B. (2015). Joint-multiple family linkage analysis predicts within-family variation better than single-family analysis of the maize nested association mapping population. Heredity, 114(6):552–563.

Parisseaux, B. and Bernardo, R. (2004). In silico mapping of quantitative trait loci in maize. Theor. Appl. Genet., 109(3):508–514.

Piepho, H.-P. and Pillen, K. (2004). Mixed modelling for QTL × environment interaction analysis. Euphytica, 137(1):147–153.

Poland, J. A., Bradbury, P. J., Buckler, E. S., and Nelson, R. J. (2011). Genome-wide nested association mapping of quantitative resistance to northern leaf blight in maize. Proc Natl Acad Sci, 108(17):6893–6898.

Saade, S., Maurer, A., Shahid, M., Oakey, H., Schmöckel, S. M., Negrão, S., Pillen, K., and Tester, M. (2016). Yield-related salinity tolerance traits identified in a nested association mapping (NAM) population of wild barley. Sci. Rep., 6:32586.

Salvi, S., Tuberosa, R., Chiapparino, E., Maccaferri, M., Veillet, S., van Beuningen, L., Isaac, P., Edwards, K., and Phillips, R. L. (2002). Toward positional cloning of Vgt1, a QTL controlling the transition from the vegetative to the reproductive phase in maize. Plant Mol. Biol., 48(5-6):601–613.

Utz, H. F., Melchinger, A. E., and Schön, C. C. (2000). Bias and sampling error of the estimated proportion of genotypic variance explained by quantitative trait loci determined from experimental data in maize using cross validation and validation with independent samples. Genetics, 154(4):1839–1849.

van Eeuwijk, F. A., Malosetti, M., and Boer, M. P. (2007). Modelling the genetic basis of response curves underlying genotype × environment interaction. In Scale and Complexity in Plant Systems Research: Gene Plant-Crop Relations, number 21, pages 115–126. Springer.

Verbyla, A. P., George, A. W., Cavanagh, C., and Verbyla, K. L. (2014). Whole-genome QTL analysis for MAGIC. Theor. Appl. Genet., 127(8):1753–1770.

Würschum, T. (2012). Mapping qtl for agronomic traits in breeding populations. Theor. Appl. Genet., 125(2):201–210.

Xavier, A., Xu, S., Muir, W. M., and Rainey, K. M. (2015). NAM: association studies in multiple populations. Bioinformatics, 31(23):3862–3864.

