## Supplementary material for "The usefulness of multi-parent multi-environment QTL analysis: an illustration in different NAM populations"

February 3, 2020

#### S1: Heritability computation

We computed the trait heritabilities on a line mean basis within the $j^{th}$ cross using the formula 3 from Hung et al. (2012)

$$h^2 = \frac{\sigma^2_{g(cr_j)}}{\sigma^2_{g(cr_j)} + \frac{\sigma^2_{gxE(cr_j)}}{N_{env}} + \frac{\sigma^2_{e(cr_j)}}{N_{env} * N_{rep}}} \quad (1)$$

##### EU-NAM data

The model used for the computation of the heritabilities was the following:

$$\begin{aligned} y_{ijklm} = & \mu + env_k + rep_{l(k)} + block_{m(lk)} + cross_j + \\ & G_i(cross_j) + G_i(cross_j) * env_k + e_{ijklm} \end{aligned} \quad (2)$$

In model 2, all terms were random and the variance of the error term was cross-specific.

| | $\sigma^2_g$ | se($\sigma^2_g$) | $\sigma^2_{ge}$ | se($\sigma^2_{ge}$) | $h^2$(%) |
|---|---|---|---|---|---|
| D152 | 208.36 | 65.38 | 93.36 | 12.26 | 60.65 |
| EC49A | 0.11 | 93.29 | 80.07 | 63.86 | 0.04 |
| EP44 | 257.42 | 139.59 | 29.11 | 176.75 | 66.64 |
| EZ5 | 41.23 | 190.46 | 0.00 | | 19.14 |
| F03802 | 145.96 | 47.45 | 528.20 | 282.09 | 27.36 |
| F2 | 302.10 | 79.10 | 75.92 | 66.24 | 71.97 |
| F283 | 361.53 | 70.04 | 0.00 | | 77.90 |
| F64 | 222.86 | 80.69 | 100.30 | 61.39 | 60.92 |
| UH006 | 224.28 | 60.12 | 138.10 | 81.42 | 57.17 |
| UH009 | 20.12 | 37.53 | 75.42 | 62.59 | 12.70 |
| DK105 | 270.12 | 75.60 | 106.34 | 61.26 | 59.76 |

#### US-NAM data

The model used for the computation of the heritabilities was the following:

$$\begin{aligned} y_{ijklmno} =& \mu + env_k + set_{l(k)} + block_{m(lk)} + row_{n(k)} + col_{o(k)} + cross_j + \\ & G_i(cross_j) + G_i * env_k + e_{ijklm} \end{aligned} \tag{3}$$

In model 3, all terms were random and the variance of the error term was modelled by an environmental specific autoregressive correlation in the row and columns (AR1 x AR1) (Gilmour et al., 1997). For the USNAM data we were not able to fit a model with within cross genotype by environment variance term so we used an homogenous genotype by environment variance term.

| | sigma.g | std.err | sigma.ge | std.err | heritability |
|---|---|---|---|---|---|
| B97 | 1.77 | 0.32 | 0.97 | 0.10 | 75.64 |
| CML103 | 1.69 | 0.37 | 0.97 | 0.10 | 64.63 |
| CML228 | 7.09 | 1.13 | 0.97 | 0.10 | 73.13 |
| CML247 | 7.19 | 0.94 | 0.97 | 0.10 | 85.04 |
| CML277 | 10.02 | 1.44 | 0.97 | 0.10 | 79.75 |
| CML322 | 3.57 | 0.54 | 0.97 | 0.10 | 82.26 |
| CML333 | 4.18 | 0.62 | 0.97 | 0.10 | 80.23 |
| CML52 | 6.20 | 0.83 | 0.97 | 0.10 | 86.47 |
| CML69 | 2.13 | 0.47 | 0.97 | 0.10 | 61.21 |
| Hp301 | 3.24 | 0.44 | 0.97 | 0.10 | 86.98 |
| IL14H | 4.32 | 0.54 | 0.97 | 0.10 | 89.91 |
| Ki11 | 6.50 | 1.06 | 0.97 | 0.10 | 70.35 |
| Ki3 | 3.79 | 0.64 | 0.97 | 0.10 | 86.53 |
| Ky21 | 1.64 | 0.32 | 0.97 | 0.10 | 71.62 |
| M162W | 3.09 | 0.50 | 0.97 | 0.10 | 78.33 |
| M37W | 3.09 | 0.53 | 0.97 | 0.10 | 73.31 |
| Mo18W | 4.84 | 0.76 | 0.97 | 0.10 | 76.34 |
| MS71 | 2.22 | 0.32 | 0.97 | 0.10 | 82.07 |
| NC350 | 2.77 | 0.51 | 0.97 | 0.10 | 70.48 |
| NC358 | 1.67 | 0.33 | 0.97 | 0.10 | 73.09 |
| Oh43 | 1.73 | 0.33 | 0.97 | 0.10 | 72.38 |
| Oh7B | 2.51 | 0.42 | 0.97 | 0.10 | 80.19 |
| P39 | 5.47 | 0.68 | 0.97 | 0.10 | 91.86 |
| Tx303 | 2.89 | 0.50 | 0.97 | 0.10 | 74.29 |
| Tzi8 | 4.60 | 0.68 | 0.97 | 0.10 | 85.03 |

### S2: List of detected QTLs

#### EU-NAM data

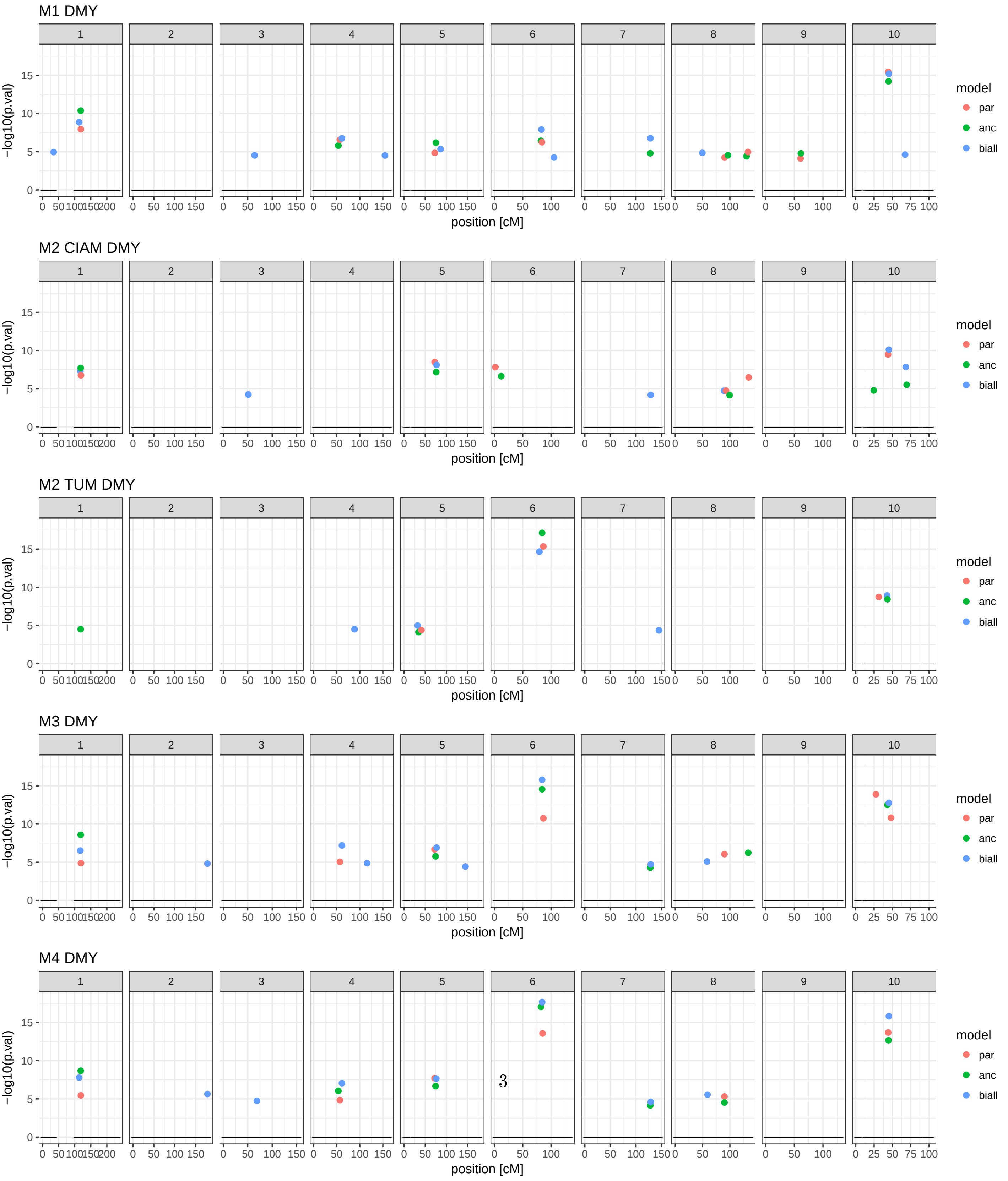


Figure 1: List of QTLs

#### EU-NAM M1

| | n | Mk. names | chr | pos [cM] | -log10(pval) |
|---|---|---|---|---|---|
| parental | | | | | |
| | 1 | PZE.101146834 | 1 | 119.1 | 8 |
| | 2 | PZE.104027223 | 4 | 56.8 | 6.6 |
| | 3 | PZE.105054186 | 5 | 72.4 | 4.8 |
| | 4 | PZE.106101027 | 6 | 83.9 | 6.3 |
| | 5 | PZE.108099840 | 8 | 90.2 | 4.2 |
| | 6 | PZE.108131921 | 8 | 133.1 | 5 |
| | 7 | PZE.109054632 | 9 | 60.9 | 4.1 |
| | 8 | PZE.110048720 | 10 | 44.3 | 15.5 |
| ancestral | | | | | |
| | 1 | PZE.101144216 | 1 | 118.6 | 10.4 |
| | 2 | PZE.104029507 | 4 | 53.5 | 5.8 |
| | 3 | PZE.105068880 | 5 | 75.2 | 6.2 |
| | 4 | PZE.106098066 | 6 | 82.1 | 6.4 |
| | 5 | PZE.107128534 | 7 | 128 | 4.8 |
| | 6 | PZE.108109731 | 8 | 96.4 | 4.5 |
| | 7 | PZE.108131479 | 8 | 130.4 | 4.4 |
| | 8 | PZE.109058296 | 9 | 61.5 | 4.8 |
| | 9 | PZE.110049068 | 10 | 44.7 | 14.2 |
| bi-allelic | | | | | |
| | 1 | PZE.101024519 | 1 | 34.3 | 5 |
| | 2 | PZE.101143233 | 1 | 113.9 | 8.9 |
| | 3 | PZE.103096063 | 3 | 64 | 4.5 |
| | 4 | PZE.104052802 | 4 | 61.2 | 6.8 |
| | 5 | PZE.104153023 | 4 | 154.5 | 4.5 |
| | 6 | PZE.105103875 | 5 | 86.4 | 5.4 |
| | 7 | PZE.106097991 | 6 | 83 | 7.9 |
| | 8 | PZE.106114241 | 6 | 105.3 | 4.3 |
| | 9 | PZE.107128336 | 7 | 128.3 | 6.8 |
| | 10 | PZE.108027746 | 8 | 49.5 | 4.9 |
| | 11 | PZE.110049474 | 10 | 45.2 | 15.2 |
| | 12 | PZE.110086343 | 10 | 67.3 | 4.6 |

#### EU-NAM M2 CIAM

| | n | Mk. names | chr | pos [cM] | -log10(pval) |
|---|---|---|---|---|---|
| parental | | | | | |
| | 1 | PZE.101147104 | 1 | 119.4 | 6.8 |
| | 2 | PZE.105054186 | 5 | 72.4 | 8.5 |
| | 3 | PZA00606.3 | 6 | 1.6 | 7.8 |
| | 4 | PZE.108104106 | 8 | 92.9 | 4.7 |
| | 5 | PZE.108133621 | 8 | 134.5 | 6.5 |
| | 6 | PZE.110049572 | 10 | 44.1 | 9.5 |
| ancestral | | | | | |
| | 1 | PZE.101144216 | 1 | 118.6 | 7.7 |
| | 2 | PZE.105063383 | 5 | 76 | 7.2 |
| | 3 | PZE.106009233 | 6 | 12 | 6.6 |
| | 4 | PZE.108110343 | 8 | 99.5 | 4.1 |
| | 5 | PZE.110009558 | 10 | 24.6 | 4.8 |
| | 6 | PZE.110087849 | 10 | 69.5 | 5.5 |
| bi-allelic | | | | | |
| | 1 | PZE.101144248 | 1 | 117.4 | 7.3 |
| | 2 | PZE.103072486 | 3 | 51.2 | 4.2 |
| | 3 | PZE.105074287 | 5 | 76.9 | 8.1 |
| | 4 | PZE.107128846 | 7 | 128.9 | 4.2 |
| | 5 | PZE.108099415 | 8 | 89.1 | 4.7 |
| | 6 | PZE.110049474 | 10 | 45.2 | 10.1 |
| | 7 | PZE.110088931 | 10 | 68.4 | 7.8 |

#### EU-NAM M2 TUM

| | n | Mk. names | chr | pos [cM] | -log10(pval) |
|---|---|---|---|---|---|
| parental | | | | | |
| | 1 | PZE.105019465 | 5 | 40.9 | 4.4 |
| | 2 | PZE.106102395 | 6 | 86.4 | 15.4 |
| | 3 | PZE.110013764 | 10 | 31.4 | 8.7 |
| ancestral | | | | | |
| | 1 | PZE.101144216 | 1 | 118.6 | 4.5 |
| | 2 | PZE.105017551 | 5 | 34.3 | 4.1 |
| | 3 | PZE.106101278 | 6 | 84.2 | 17.1 |
| | 4 | PZE.110049040 | 10 | 43.2 | 8.4 |
| bi-allelic | | | | | |
| | 1 | PZE.104094429 | 4 | 88.6 | 4.5 |
| | 2 | PZE.105017975 | 5 | 32 | 5 |
| | 3 | PZE.106095383 | 6 | 79.3 | 14.7 |
| | 4 | PZE.107136612 | 7 | 145 | 4.4 |
| | 5 | PZE.110049406 | 10 | 42.8 | 8.9 |

#### EU-NAM M3

| | n | Mk. names | chr | pos [cM] | -log10(pval) |
|---|---|---|---|---|---|
| parental | | | | | |
| | 1 | PZE.101147104 | 1 | 119.4 | 4.9 |
| | 2 | PZE.104027223 | 4 | 56.8 | 5.1 |
| | 3 | PZE.105062183 | 5 | 72.2 | 6.7 |
| | 4 | PZE.106102395 | 6 | 86.4 | 10.8 |
| | 5 | PZE.108099425 | 8 | 90 | 6.1 |
| | 6 | PZE.110010098 | 10 | 27.5 | 13.9 |
| | 7 | PZE.110049922 | 10 | 48.1 | 10.8 |
| ancestral | | | | | |
| | 1 | PZE.101144216 | 1 | 118.6 | 8.6 |
| | 2 | PZE.105065789 | 5 | 74.5 | 5.8 |
| | 3 | PZE.106101278 | 6 | 84.2 | 14.6 |
| | 4 | PZE.107128534 | 7 | 128 | 4.3 |
| | 5 | PZE.108133100 | 8 | 133.6 | 6.2 |
| | 6 | PZE.110049040 | 10 | 43.2 | 12.5 |
| bi-allelic | | | | | |
| | 1 | PZE.101144248 | 1 | 117.4 | 6.5 |
| | 2 | PZE.102192367 | 2 | 178.2 | 4.8 |
| | 3 | PZE.104052802 | 4 | 61.2 | 7.2 |
| | 4 | PZE.104110016 | 4 | 115.6 | 4.9 |
| | 5 | PZE.105074287 | 5 | 76.9 | 6.9 |
| | 6 | PZE.105160757 | 5 | 145.1 | 4.4 |
| | 7 | PZE.106101278 | 6 | 84.2 | 15.8 |
| | 8 | PZE.107128846 | 7 | 128.9 | 4.7 |
| | 9 | PZE.108057679 | 8 | 58.2 | 5.1 |
| | 10 | PZE.110049474 | 10 | 45.2 | 12.8 |

#### EU-NAM M4

| | n | Mk. names | chr | pos [cM] | -log10(pval) |
|---|---|---|---|---|---|
| parental | | | | | |
| | 1 | PZE.101146834 | 1 | 119.1 | 5.5 |
| | 2 | PZE.104027223 | 4 | 56.8 | 4.9 |
| | 3 | PZE.105062183 | 5 | 72.2 | 7.7 |
| | 4 | PZE.106099144 | 6 | 85 | 13.6 |
| | 5 | PZE.108099425 | 8 | 90 | 5.3 |
| | 6 | PZE.110048720 | 10 | 44.3 | 13.7 |
| ancestral | | | | | |
| | 1 | PZE.101144216 | 1 | 118.6 | 8.7 |
| | 2 | PZE.104029507 | 4 | 53.5 | 6.1 |
| | 3 | PZE.105065789 | 5 | 74.5 | 6.7 |
| | 4 | PZE.106098066 | 6 | 82.1 | 17 |
| | 5 | PZE.107128534 | 7 | 128 | 4.1 |
| | 6 | PZE.108099425 | 8 | 90 | 4.5 |
| | 7 | PZE.110049068 | 10 | 44.7 | 12.7 |
| bi-allelic | | | | | |
| | 1 | PZE.101143233 | 1 | 113.9 | 7.8 |
| | 2 | PZE.102192367 | 2 | 178.2 | 5.6 |
| | 3 | PZE.103106593 | 3 | 68.7 | 4.8 |
| | 4 | PZE.104052802 | 4 | 61.2 | 7.1 |
| | 5 | PZE.105063758 | 5 | 76.1 | 7.7 |
| | 6 | PZE.106101278 | 6 | 84.2 | 17.7 |
| | 7 | PZE.107128846 | 7 | 128.9 | 4.6 |
| | 8 | PZE.108058577 | 8 | 59.1 | 5.6 |
| | 9 | PZE.110049474 | 10 | 45.2 | 15.8 |

#### US-NAM data

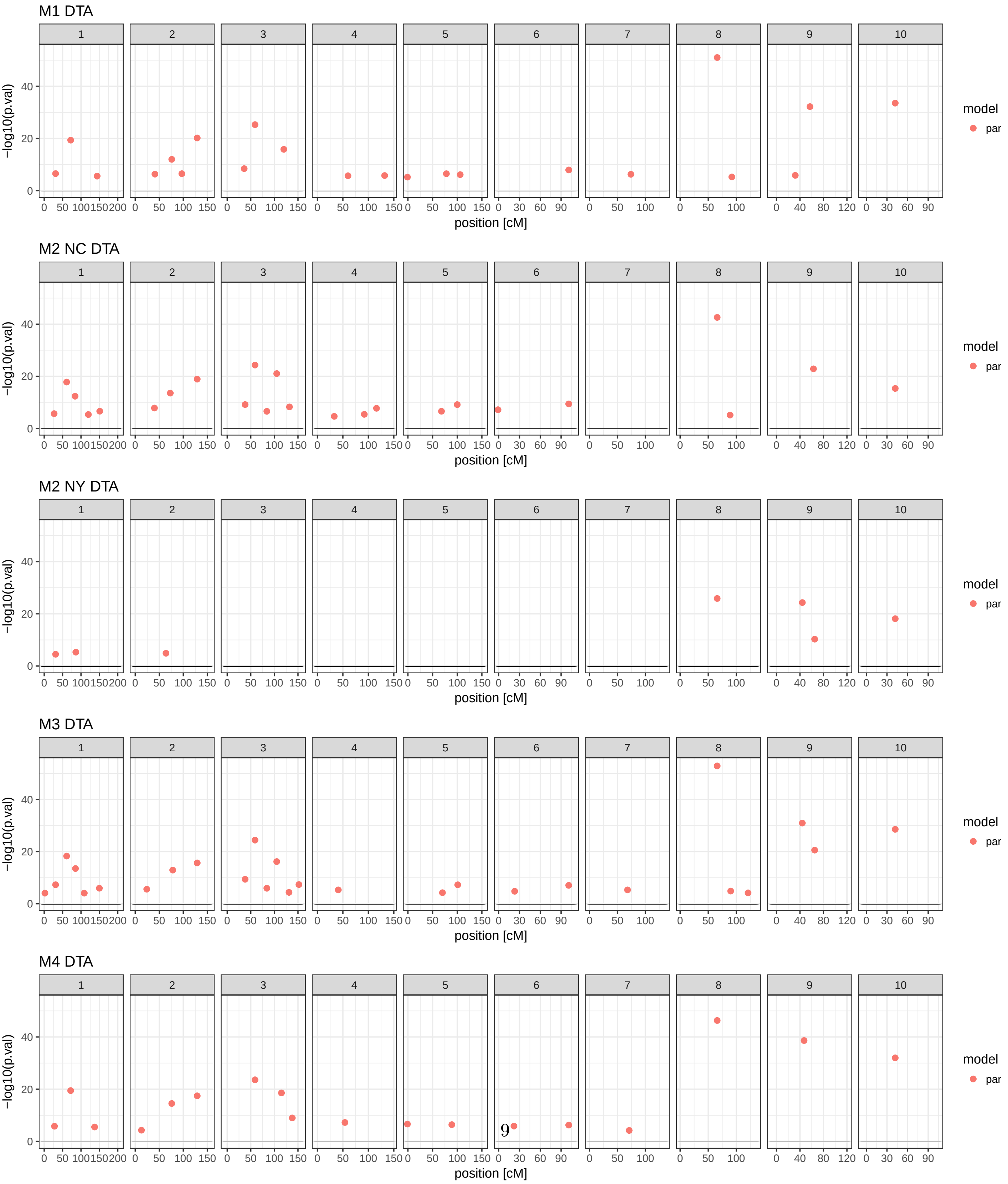


Figure 2: List of QTLs

#### US-NAM M1

| | n | Mk. names | chr | pos [cM] | -log10(pval) |
|---|---|---|---|---|---|
| parental | | | | | |
| | 1 | m36 | 1 | 31 | 6.5 |
| | 2 | m77 | 1 | 72 | 19.4 |
| | 3 | m149 | 1 | 144 | 5.6 |
| | 4 | m255 | 2 | 41 | 6.3 |
| | 5 | m290 | 2 | 76 | 12 |
| | 6 | m311 | 2 | 97 | 6.5 |
| | 7 | m343 | 2 | 129 | 20.2 |
| | 8 | m413 | 3 | 36 | 8.5 |
| | 9 | m436 | 3 | 59 | 25.3 |
| | 10 | m497 | 3 | 120 | 15.8 |
| | 11 | m599 | 4 | 60 | 5.7 |
| | 12 | m671 | 4 | 132 | 5.8 |
| | 13 | m689 | 5 | -1 | 5.2 |
| | 14 | m768 | 5 | 78 | 6.5 |
| | 15 | m796 | 5 | 106 | 6.2 |
| | 16 | m947 | 6 | 101 | 7.9 |
| | 17 | m1032 | 7 | 74 | 6.3 |
| | 18 | m1162 | 8 | 66 | 51 |
| | 19 | m1188 | 8 | 92 | 5.3 |
| | 20 | m1275 | 9 | 32 | 5.9 |
| | 21 | m1300 | 9 | 57 | 32.2 |
| | 22 | m1414 | 10 | 42 | 33.5 |

#### US-NAM M2 New York

| | n | Mk. names | chr | pos [cM] | -log10(pval) |
|---|---|---|---|---|---|
| parental | | | | | |
| | 1 | m32 | 1 | 27 | 5.7 |
| | 2 | m66 | 1 | 61 | 17.8 |
| | 3 | m89 | 1 | 84 | 12.3 |
| | 4 | m125 | 1 | 120 | 5.3 |
| | 5 | m156 | 1 | 151 | 6.6 |
| | 6 | m254 | 2 | 40 | 7.8 |
| | 7 | m287 | 2 | 73 | 13.5 |
| | 8 | m343 | 2 | 129 | 18.9 |
| | 9 | m415 | 3 | 38 | 9.1 |
| | 10 | m436 | 3 | 59 | 24.3 |
| | 11 | m461 | 3 | 84 | 6.5 |
| | 12 | m482 | 3 | 105 | 21 |
| | 13 | m509 | 3 | 132 | 8.2 |
| | 14 | m572 | 4 | 33 | 4.6 |
| | 15 | m631 | 4 | 92 | 5.4 |
| | 16 | m655 | 4 | 116 | 7.7 |
| | 17 | m758 | 5 | 68 | 6.6 |
| | 18 | m790 | 5 | 100 | 9.1 |
| | 19 | m845 | 6 | -1 | 7.2 |
| | 20 | m947 | 6 | 101 | 9.4 |
| | 21 | m1162 | 8 | 66 | 42.6 |
| | 22 | m1185 | 8 | 89 | 5.1 |
| | 23 | m1306 | 9 | 63 | 22.9 |
| | 24 | m1414 | 10 | 42 | 15.3 |

#### US-NAM M2 North Carolina

| | n | Mk. names | chr | pos [cM] | -log10(pval) |
|---|---|---|---|---|---|
| parental | | | | | |
| | 1 | m36 | 1 | 31 | 4.5 |
| | 2 | m91 | 1 | 86 | 5.3 |
| | 3 | m278 | 2 | 64 | 4.9 |
| | 4 | m1162 | 8 | 66 | 25.9 |
| | 5 | m1287 | 9 | 44 | 24.3 |
| | 6 | m1308 | 9 | 65 | 10.3 |
| | 7 | m1414 | 10 | 42 | 18.2 |

#### US-NAM M3

| | n | Mk. names | chr | pos [cM] | -log10(pval) |
|---|---|---|---|---|---|
| parental | | | | | |
| | 1 | m7 | 1 | 2 | 4.1 |
| | 2 | m36 | 1 | 31 | 7.3 |
| | 3 | m66 | 1 | 61 | 18.3 |
| | 4 | m90 | 1 | 85 | 13.5 |
| | 5 | m114 | 1 | 109 | 4.1 |
| | 6 | m155 | 1 | 150 | 6 |
| | 7 | m238 | 2 | 24 | 5.6 |
| | 8 | m292 | 2 | 78 | 12.9 |
| | 9 | m343 | 2 | 129 | 15.7 |
| | 10 | m415 | 3 | 38 | 9.4 |
| | 11 | m436 | 3 | 59 | 24.4 |
| | 12 | m461 | 3 | 84 | 5.9 |
| | 13 | m482 | 3 | 105 | 16.2 |
| | 14 | m508 | 3 | 131 | 4.4 |
| | 15 | m529 | 3 | 152 | 7.4 |
| | 16 | m580 | 4 | 41 | 5.3 |
| | 17 | m760 | 5 | 70 | 4.2 |
| | 18 | m791 | 5 | 101 | 7.3 |
| | 19 | m869 | 6 | 23 | 4.8 |
| | 20 | m947 | 6 | 101 | 7.1 |
| | 21 | m1026 | 7 | 68 | 5.3 |
| | 22 | m1162 | 8 | 66 | 52.9 |
| | 23 | m1186 | 8 | 90 | 4.9 |
| | 24 | m1217 | 8 | 121 | 4.2 |
| | 25 | m1287 | 9 | 44 | 31 |
| | 26 | m1308 | 9 | 65 | 20.6 |
| | 27 | m1414 | 10 | 42 | 28.6 |

#### US-NAM M4

| | n | Mk. names | chr | pos [cM] | -log10(pval) |
|---|---|---|---|---|---|
| parental | | | | | |
| | 1 | m33 | 1 | 28 | 5.8 |
| | 2 | m77 | 1 | 72 | 19.5 |
| | 3 | m142 | 1 | 137 | 5.5 |
| | 4 | m227 | 2 | 13 | 4.3 |
| | 5 | m290 | 2 | 76 | 14.5 |
| | 6 | m343 | 2 | 129 | 17.5 |
| | 7 | m436 | 3 | 59 | 23.6 |
| | 8 | m492 | 3 | 115 | 18.6 |
| | 9 | m515 | 3 | 138 | 9 |
| | 10 | m593 | 4 | 54 | 7.2 |
| | 11 | m689 | 5 | -1 | 6.6 |
| | 12 | m779 | 5 | 89 | 6.4 |
| | 13 | m868 | 6 | 22 | 5.9 |
| | 14 | m947 | 6 | 101 | 6.3 |
| | 15 | m1029 | 7 | 71 | 4.2 |
| | 16 | m1162 | 8 | 66 | 46.3 |
| | 17 | m1290 | 9 | 47 | 38.6 |
| | 18 | m1414 | 10 | 42 | 32 |

### S3: QTL additive effects allelic series

#### EU-NAM chr 6 82.1 cM

Table 1: QTL additive effects and standard deviations

| | $\beta_{M1}$ | $\beta_{M4-E1}$ | $\beta_{M4-E2}$ | sd($\beta_{M1}$) | sd($\beta_{M4-E1}$) | sd($\beta_{M4-E2}$) | $\beta$/sd($\beta$)(M1) | $\beta$/sd($\beta$)(M4-E1) | $\beta$/sd($\beta$)(M4-E2) |
|---|---|---|---|---|---|---|---|---|---|
| UH007 | 0.00 | 0.00 | 0.00 | 0.00 | 0.00 | 0.00 | | | |
| EZ5 | 0.00 | 0.00 | 0.00 | 0.00 | 0.00 | 0.00 | | | |
| UH009 | 0.00 | 0.00 | 0.00 | 0.00 | 0.00 | 0.00 | | | |
| D152 | -4.07 | -1.19 | -7.20 | 0.71 | 0.83 | 0.83 | -5.75 | -1.42 | -8.63 |
| EC49A | -4.07 | -1.19 | -7.20 | 0.71 | 0.83 | 0.83 | -5.75 | -1.42 | -8.63 |
| F03802 | -4.07 | -1.19 | -7.20 | 0.71 | 0.83 | 0.83 | -5.75 | -1.42 | -8.63 |
| F2 | -4.07 | -1.19 | -7.20 | 0.71 | 0.83 | 0.83 | -5.75 | -1.42 | -8.63 |
| F283 | -4.07 | -1.19 | -7.20 | 0.71 | 0.83 | 0.83 | -5.75 | -1.42 | -8.63 |
| UH006 | -4.07 | -1.19 | -7.20 | 0.71 | 0.83 | 0.83 | -5.75 | -1.42 | -8.63 |
| DK105 | -4.07 | -1.19 | -7.20 | 0.71 | 0.83 | 0.83 | -5.75 | -1.42 | -8.63 |
| F64 | -6.13 | 0.52 | -12.81 | 2.13 | 2.40 | 2.40 | -2.88 | 0.22 | -5.34 |
| EP44 | -0.94 | 5.65 | 10.19 | 5.72 | 5.63 | 5.63 | -0.17 | 1.00 | 1.81 |

#### US-NAM chr 8 66 cM

Table 2: QTL additive effects and standard deviations

| | $\beta_{M1}$ | $\beta_{M4-E1}$ | $\beta_{M4-E2}$ | sd($\beta_{M1}$) | sd($\beta_{M4-E1}$) | sd($\beta_{M4-E2}$) | $\beta$/sd($\beta$)(M1) | $\beta$/sd($\beta$)(M4-E1) | $\beta$/sd($\beta$)(M4-E2) |
|---|---|---|---|---|---|---|---|---|---|
| B73 | 0.00 | 0.00 | 0.00 | 0.00 | 0.00 | 0.00 | | | |
| B97 | 0.04 | 0.09 | 0.08 | 0.14 | 0.14 | 0.14 | 0.27 | 0.66 | 0.56 |
| CML103 | 0.29 | 0.55 | 0.21 | 0.16 | 0.16 | 0.16 | 1.83 | 3.46 | 1.31 |
| CML228 | 0.36 | 0.43 | 0.45 | 0.22 | 0.22 | 0.22 | 1.61 | 1.92 | 2.03 |
| CML247 | 0.54 | 0.45 | 0.55 | 0.17 | 0.20 | 0.20 | 3.12 | 2.28 | 2.83 |
| CML277 | -0.05 | 0.26 | -0.17 | 0.22 | 0.18 | 0.18 | -0.20 | 1.44 | -0.94 |
| CML322 | 0.41 | 0.68 | 0.60 | 0.16 | 0.18 | 0.18 | 2.62 | 3.91 | 3.45 |
| CML333 | 0.60 | 0.69 | 0.62 | 0.17 | 0.19 | 0.19 | 3.55 | 3.65 | 3.30 |
| CML52 | 0.59 | 0.47 | 0.52 | 0.22 | 0.19 | 0.19 | 2.65 | 2.49 | 2.73 |
| CML69 | 0.61 | 0.67 | 0.64 | 0.16 | 0.16 | 0.16 | 3.70 | 4.16 | 3.98 |
| Hp301 | 0.08 | 0.20 | 0.19 | 0.14 | 0.14 | 0.14 | 0.54 | 1.43 | 1.31 |
| IL14H | -1.09 | -0.82 | -1.32 | 0.14 | 0.14 | 0.14 | -7.56 | -5.80 | -9.35 |
| Ki11 | -0.11 | -0.06 | -0.52 | 0.19 | 0.20 | 0.20 | -0.54 | -0.29 | -2.61 |
| Ki3 | 0.17 | 0.44 | -0.07 | 0.20 | 0.29 | 0.29 | 0.86 | 1.51 | -0.23 |
| Ky21 | 0.38 | 0.32 | 0.46 | 0.12 | 0.13 | 0.13 | 3.09 | 2.43 | 3.49 |
| M162W | 0.03 | -0.03 | -0.11 | 0.13 | 0.16 | 0.16 | 0.25 | -0.17 | -0.72 |
| M37W | 0.20 | 0.17 | 0.00 | 0.16 | 0.15 | 0.15 | 1.24 | 1.14 | 0.02 |
| Mo18W | 0.62 | 0.87 | 0.53 | 0.18 | 0.20 | 0.20 | 3.54 | 4.32 | 2.64 |
| MS71 | -1.06 | -0.54 | -1.17 | 0.14 | 0.13 | 0.13 | -7.54 | -4.13 | -8.99 |
| NC350 | -0.27 | 0.56 | -0.20 | 0.18 | 0.18 | 0.18 | -1.45 | 3.11 | -1.10 |
| NC358 | 0.04 | 0.40 | -0.05 | 0.12 | 0.14 | 0.14 | 0.32 | 2.81 | -0.34 |
| Oh43 | -0.02 | -0.07 | 0.12 | 0.13 | 0.14 | 0.14 | -0.14 | -0.50 | 0.81 |
| Oh7B | 0.15 | 0.22 | 0.20 | 0.15 | 0.16 | 0.16 | 1.01 | 1.33 | 1.24 |
| P39 | -1.32 | -0.99 | -1.56 | 0.18 | 0.18 | 0.18 | -7.46 | -5.48 | -8.69 |
| Tx303 | 0.48 | 0.64 | 0.44 | 0.17 | 0.20 | 0.20 | 2.87 | 3.24 | 2.23 |
| Tzi8 | -0.11 | -0.23 | -0.44 | 0.15 | 0.20 | 0.20 | -0.74 | -1.17 | -2.18 |

#### US-NAM chr 9 47 cM

Table 3: QTL additive effects and standard deviations

| | $\beta_{M1}$ | $\beta_{M4-E1}$ | $\beta_{M4-E2}$ | sd($\beta_{M1}$) | sd($\beta_{M4-E1}$) | sd($\beta_{M4-E2}$) | $\beta$/sd($\beta$)(M1) | $\beta$/sd($\beta$)(M4-E1) | $\beta$/sd($\beta$)(M4-E2) |
|---|---|---|---|---|---|---|---|---|---|
| B73 | 0.00 | 0.00 | 0.00 | 0.00 | 0.00 | 0.00 | | | |
| B97 | -0.05 | 0.01 | 0.02 | 0.12 | 0.13 | 0.13 | -0.40 | 0.06 | 0.18 |
| CML103 | 0.47 | 0.25 | 0.69 | 0.13 | 0.16 | 0.16 | 3.62 | 1.57 | 4.44 |
| CML228 | 0.93 | 0.67 | 1.20 | 0.22 | 0.23 | 0.23 | 4.26 | 2.95 | 5.30 |
| CML247 | 0.20 | -0.29 | 0.71 | 0.16 | 0.19 | 0.19 | 1.25 | -1.52 | 3.70 |
| CML277 | 1.43 | 0.78 | 1.94 | 0.21 | 0.18 | 0.18 | 6.85 | 4.41 | 11.00 |
| CML322 | 0.53 | 0.30 | 0.79 | 0.14 | 0.17 | 0.17 | 3.73 | 1.79 | 4.65 |
| CML333 | 0.70 | 0.26 | 1.08 | 0.16 | 0.19 | 0.19 | 4.39 | 1.39 | 5.71 |
| CML52 | 0.48 | 0.28 | 0.82 | 0.21 | 0.18 | 0.18 | 2.30 | 1.55 | 4.62 |
| CML69 | 0.03 | -0.36 | 0.18 | 0.14 | 0.15 | 0.15 | 0.19 | -2.42 | 1.23 |
| Hp301 | 0.46 | 0.51 | 0.41 | 0.13 | 0.14 | 0.14 | 3.42 | 3.56 | 2.81 |
| IL14H | -0.05 | 0.15 | 0.18 | 0.14 | 0.14 | 0.14 | -0.37 | 1.07 | 1.30 |
| Ki11 | 1.10 | 0.28 | 1.85 | 0.20 | 0.21 | 0.21 | 5.43 | 1.33 | 8.70 |
| Ki3 | 0.12 | -0.19 | 0.20 | 0.18 | 0.27 | 0.27 | 0.65 | -0.69 | 0.76 |
| Ky21 | 0.23 | 0.29 | 0.18 | 0.10 | 0.13 | 0.13 | 2.23 | 2.20 | 1.35 |
| M162W | 0.36 | -0.11 | 0.65 | 0.14 | 0.18 | 0.18 | 2.56 | -0.62 | 3.66 |
| M37W | -0.04 | -0.02 | 0.15 | 0.15 | 0.16 | 0.16 | -0.24 | -0.13 | 0.96 |
| Mo18W | 0.16 | -0.36 | 0.60 | 0.17 | 0.21 | 0.21 | 0.93 | -1.72 | 2.82 |
| MS71 | 0.31 | 0.04 | 0.46 | 0.13 | 0.13 | 0.13 | 2.42 | 0.32 | 3.58 |
| NC350 | -0.48 | -0.57 | -0.09 | 0.15 | 0.18 | 0.18 | -3.28 | -3.17 | -0.51 |
| NC358 | -0.07 | -0.29 | 0.36 | 0.12 | 0.15 | 0.15 | -0.54 | -2.01 | 2.47 |
| Oh43 | 0.14 | 0.06 | 0.22 | 0.13 | 0.14 | 0.14 | 1.07 | 0.45 | 1.51 |
| Oh7B | 0.28 | 0.38 | 0.36 | 0.14 | 0.17 | 0.17 | 2.03 | 2.20 | 2.08 |
| P39 | 0.28 | 0.55 | 0.50 | 0.17 | 0.19 | 0.19 | 1.62 | 2.85 | 2.57 |
| Tx303 | 0.21 | -0.09 | 0.51 | 0.15 | 0.20 | 0.20 | 1.42 | -0.42 | 2.52 |
| Tzi8 | 0.72 | 0.45 | 0.92 | 0.13 | 0.18 | 0.18 | 5.42 | 2.45 | 5.02 |

#### US-NAM chr 10 42 cM

Table 4: QTL additive effects and standard deviations

| | $\beta_{M1}$ | $\beta_{M4-E1}$ | $\beta_{M4-E2}$ | sd($\beta_{M1}$) | sd($\beta_{M4-E1}$) | sd($\beta_{M4-E2}$) | $\beta$/sd($\beta$)(M1) | $\beta$/sd($\beta$)(M4-E1) | $\beta$/sd($\beta$)(M4-E2) |
|---|---|---|---|---|---|---|---|---|---|
| B73 | 0.00 | 0.00 | 0.00 | 0.00 | 0.00 | 0.00 | | | |
| B97 | -0.05 | 0.04 | -0.20 | 0.13 | 0.13 | 0.13 | -0.42 | 0.30 | -1.49 |
| CML103 | -0.21 | -0.20 | -0.36 | 0.13 | 0.16 | 0.16 | -1.61 | -1.21 | -2.21 |
| CML228 | 1.34 | 0.66 | 2.10 | 0.21 | 0.22 | 0.22 | 6.27 | 2.98 | 9.41 |
| CML247 | -0.14 | 0.01 | -0.47 | 0.16 | 0.19 | 0.19 | -0.87 | 0.07 | -2.43 |
| CML277 | 2.03 | 1.39 | 2.99 | 0.22 | 0.18 | 0.18 | 9.22 | 7.54 | 16.26 |
| CML322 | -0.45 | -0.26 | -0.50 | 0.14 | 0.17 | 0.17 | -3.22 | -1.57 | -2.97 |
| CML333 | 0.16 | -0.12 | 0.14 | 0.16 | 0.19 | 0.19 | 1.02 | -0.66 | 0.74 |
| CML52 | 0.29 | 0.41 | 0.22 | 0.20 | 0.19 | 0.19 | 1.45 | 2.20 | 1.21 |
| CML69 | -0.12 | 0.16 | -0.26 | 0.14 | 0.16 | 0.16 | -0.85 | 1.04 | -1.63 |
| Hp301 | 0.20 | -0.01 | 0.41 | 0.14 | 0.15 | 0.15 | 1.44 | -0.04 | 2.79 |
| IL14H | -0.05 | 0.50 | -0.02 | 0.13 | 0.14 | 0.14 | -0.37 | 3.62 | -0.15 |
| Ki11 | 1.48 | 1.09 | 1.76 | 0.19 | 0.20 | 0.20 | 7.65 | 5.37 | 8.69 |
| Ki3 | -0.43 | -0.55 | -0.44 | 0.18 | 0.28 | 0.28 | -2.38 | -1.97 | -1.57 |
| Ky21 | 0.23 | 0.24 | 0.33 | 0.10 | 0.13 | 0.13 | 2.22 | 1.76 | 2.44 |
| M162W | -0.08 | -0.03 | -0.37 | 0.14 | 0.18 | 0.18 | -0.61 | -0.20 | -2.12 |
| M37W | -0.17 | -0.17 | -0.23 | 0.14 | 0.15 | 0.15 | -1.22 | -1.13 | -1.56 |
| Mo18W | -0.21 | -0.14 | -0.42 | 0.17 | 0.21 | 0.21 | -1.23 | -0.65 | -1.98 |
| MS71 | 0.17 | 0.04 | 0.26 | 0.14 | 0.14 | 0.14 | 1.21 | 0.31 | 1.86 |
| NC350 | -0.20 | -0.02 | -0.07 | 0.15 | 0.18 | 0.18 | -1.31 | -0.13 | -0.43 |
| NC358 | 0.22 | 0.27 | 0.26 | 0.12 | 0.14 | 0.14 | 1.87 | 1.90 | 1.81 |
| Oh43 | -0.12 | -0.05 | -0.15 | 0.13 | 0.15 | 0.15 | -0.91 | -0.33 | -1.01 |
| Oh7B | -0.01 | 0.18 | 0.09 | 0.13 | 0.16 | 0.16 | -0.07 | 1.11 | 0.53 |
| P39 | -0.15 | 0.00 | -0.16 | 0.16 | 0.18 | 0.18 | -0.90 | 0.02 | -0.89 |
| Tx303 | -0.17 | -0.04 | -0.15 | 0.14 | 0.20 | 0.20 | -1.19 | -0.20 | -0.75 |
| Tzi8 | -0.03 | -0.08 | -0.35 | 0.14 | 0.19 | 0.19 | -0.23 | -0.40 | -1.82 |

### S4: QTL additive effects allelic series

Comparison of the additive effect allelic series between M1, M2, M3 and M4 and two environments for two individual QTL positions. The upper panel contain the first environment result and the lower the one from the second environment.

#### EU-NAM ancestral model chr 6 82.1 cM

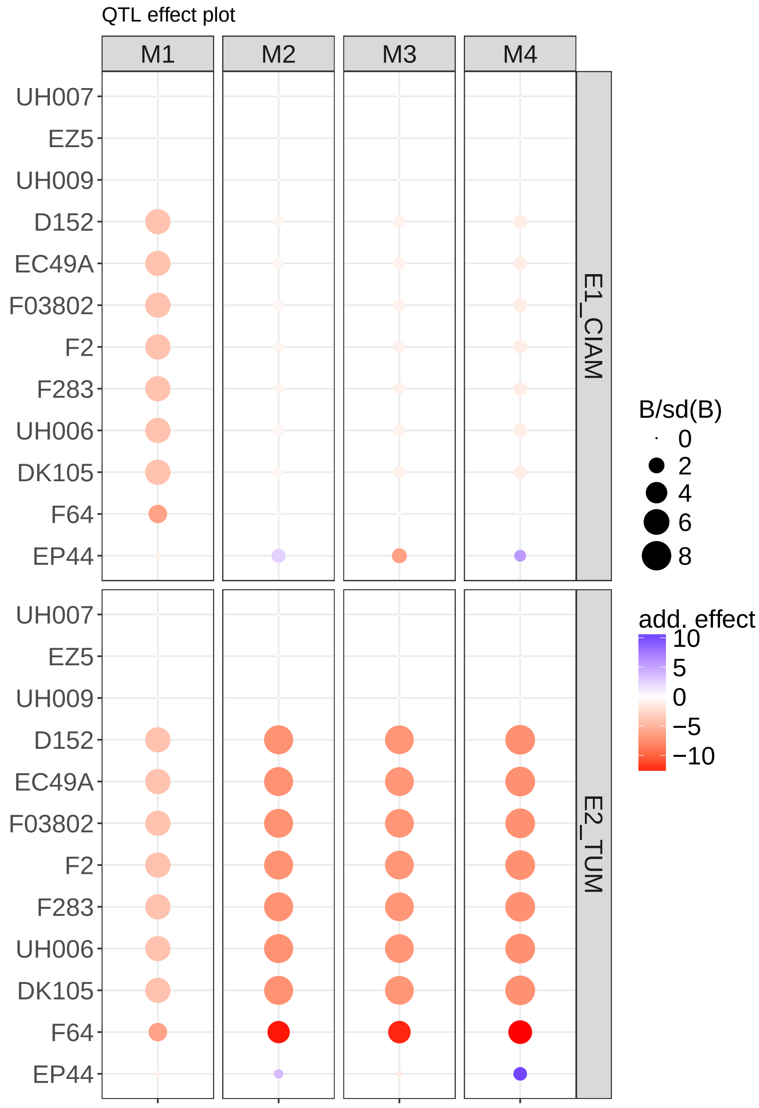


Figure 3: Comparison of the additive effect allelic series between M1, M2, M3 and M4 and two environments

#### US-NAM parental model chr 8 67 cM

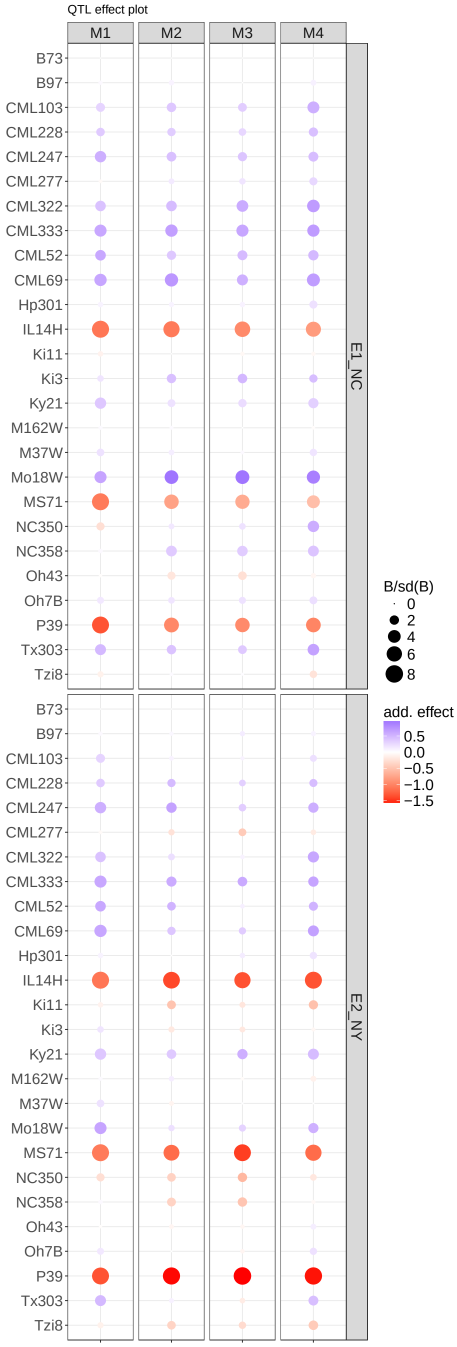


Figure 4: Comparison of the additive effect allelic series between M1, M2, M3 and M4 and two environments

### References


Gilmour, A. R., Cullis, B. R., and Verbyla, A. P. (1997). Accounting for natural and extraneous variation in the analysis of field experiments. *Journal of Agricultural, Biological, and Environmental Statistics*, pages 269–293.

Hung, H., Browne, C., Guill, K., Coles, N., Eller, M., Garcia, A., Lepak, N., Melia-Hancock, S., Oropeza-Rosas, M., Salvo, S., et al. (2012). The relationship between parental genetic or phenotypic divergence and progeny variation in the maize nested association mapping population. *Heredity*, 108(5):490–499.
